# IL-17 producing tissue-resident memory T-cells expanded during *Staphylococcus aureus* nasal colonisation provide heterologous immune protection

**DOI:** 10.64898/2026.08.31.747503

**Authors:** Clíodhna M. Daly, Seán C. Cahill, Charlotte M. Leane, Simon R. Carlile, Alanna M. Kelly, Jenny M. Mannion, Rachel M. McLoughlin

## Abstract

*Staphylococcus aureus* persistently colonises the nasal tissue (NT) of a significant proportion of the population. The long-lasting impact that asymptomatic *S. aureus* exposure has on immune memory at colonised barrier sites is incompletely understood, potentially impacting vaccine responsiveness in a pre-exposed population. Tissue resident memory (TRM) cells are long-lived T-cells which remain poised at barrier sites for localised reactivation following antigen exposure. This study demonstrates an increase in NT CD4+ and γδ+ TRM cells in response to *S. aureus* colonisation, which undergo expansion and IL-17 production upon secondary *S. aureus* exposure. Interestingly, these cells were also capable of non-specific reactivation, with IL-17+ TRM cells in *S. aureus* colonised mice enhancing protection against *K. pneumoniae* infection. Ex-vivo data suggest that non-specific CD4+ TRM cell re-activation is pro-inflammatory cytokine dependent, but antigen independent. Overall, these findings demonstrate that *S. aureus* nasal colonisation shapes long-lasting TRM cell responses in the NT, which have the potential for non-specific bystander reactivation during subsequent heterologous infection.

## Introduction

*Staphylococcus aureus* is a commonly encountered pathobiont, that asymptomatically colonises sites such as the skin, axilla, gastrointestinal tract and nasal mucosa [1]. 20-30% of healthy individuals are thought to persistently carry *S. aureus* in the anterior nares, with the remainder of individuals encountering the bacterium transiently over their lifetime [2]. *S. aureus* can also drive severe, invasive infections with high levels of fatality [3] when barrier integrity is breached, or where host immunity is compromised [4]. Treatment of *S. aureus* infections is becoming increasingly difficult as strains become progressively resistant to antibiotic treatment [5], highlighting the urgent need for a vaccine to protect against *S. aureus* infections. While significant efforts have been deployed, to-date, an efficacious vaccine against *S. aureus* remains elusive [6]. A key consideration absent from previous vaccine trials is the impact that prior exposure to this bacterium might have on shaping the host’s immune response, with significant emerging evidence suggesting that prior *S. aureus* exposure may lead to vaccine interference [6]–[9]. *S. aureus* colonisation status has been reported to influence outcomes of invasive *S. aureus* infection, with lower severity and mortality risk observed among colonised individuals that develop bacteraemia compared to non-colonised, infected individuals [10]. This suggests a potential immunological advantage associated with *S. aureus* colonisation. However, the duality of this bacterium as both a colonising organism and an infectious pathogen results in a nuanced relationship between *S. aureus* and the host immune system which is overall incompletely understood [6].

In murine models, effective clearance of *S. aureus* from the nasal tissue (NT) requires a robust T-cell response, specifically, activation of IL-17 and IL-22 producing T-cells. IL-17 acts as a key mediator of neutrophil recruitment to the nasal lumen, which is essential for *S. aureus* clearance [11],[12]. IL-22 has been shown to facilitate local antimicrobial peptide (AMP) production, as well as controlling the expression of host ligands loricrin and Cytokeratin10, which are targeted by *S. aureus* for binding at this site [13]. Human studies have demonstrated that rates of *S. aureus* nasal colonisation are higher in HIV+ individuals, where CD4+ T-cell function is compromised, once again indicating a critical role for CD4+ T-cells in *S. aureus* decolonisation [14]. Despite reports of T-cell activity in the *S. aureus* colonised NT, it remains to be established whether *S. aureus-*reactive T-cells re-enter the circulation or remain resident within the NT where they could be acting as crucial contributors to barrier immunity and protection in the nasal mucosa.

Tissue Resident Memory cells (TRM cells) are a T-cell population that do not re-enter the circulation, remaining poised at barrier sites for rapid reactivation upon secondary antigen exposure, providing localised protective immunity and immunosurveillance [15]. NT TRM cells that are expanded in response to microbial stimulation [16] or localised vaccine administration [17],[18] are retained in the nasal mucosa for rapid reactivation upon subsequent challenge. *Bordetella pertussis* and *Streptococcus pneumoniae* have both been shown to drive IL-17+ CD4+ TRM cell expansion in the nasal mucosa, which enhance neutrophil recruitment upon subsequent re-exposure to the priming bacteria [19]–[21]. Interestingly, *B. pertussis*-specific CD4+ nasal TRM cells expanded in response to vaccination have also been found to provide heterologous protection, undergoing non-specific reactivation following *Klebsiella pneumoniae* or Lipopolysaccharide (LPS) exposure [16]. These studies highlight the potential protective roles of nasal TRM cells that are expanded following bacterial exposure.

While studies to date have primarily focused on TRM cell generation following infection or vaccination, there is evidence to suggest that TRM cell phenotype and function can be influenced by colonising microbes. TRM cell heterogeneity across tissue sites is thought to be driven by location-specific factors such as the presence of other immune cell populations and varying levels of cytokines, often as a direct result of diverse microbial landscapes that make up distinct tissue microenvironments [22],[23]. In the skin Naik et al. demonstrated that colonisation with *Staphylococcus epidermidis* drives upregulation of CD4+ TRM cells which enhance barrier immunity and promote IL-17-dependent heterologous protection against invasive *Candida albicans* [24]. Intestinal TRM cell function is also thought to be shaped by the commensal microbial populations present [25]. Collectively, these findings suggest that TRM cells can be generated in response to colonising microbes that exist as part of the tissue microbiome at barrier sites, potentially contributing to protection against invasive infection by alternative pathogens.

Although acute T-cell responses to *S. aureus* colonisation have been investigated, in situ *S. aureus* memory responses at colonised barrier sites have not been fully explored. This study demonstrates that nasal colonisation with *S. aureus* leads to generation of CD4+ and γδ+ TRM cells localised within the colonised nasal mucosa. Increased IL-17 production by TRM cells enhances *K. pneumoniae* clearance from *S. aureus* colonised mice compared to non-colonised mice, indicating a protective role for *S. aureus-*primed NT TRM cells during non-specific bacterial infections. Bystander activation of *S. aureus*-primed CD4+ TRM cells during heterologous infection appears to be driven by cytokine signalling and not TCR engagement. These findings identify a strong TRM cell imprint induced by *S. aureus*, colonisation, with potential for protective effects during heterologous infection.

## Results

### *S. aureus* colonisation drives TRM cell expansion in the NT

To investigate local T-cell responses within the NT during *S. aureus* colonisation, wild-type C57BL6/JCrl mice were intranasally (i.n.) inoculated with streptomycin-resistant *S. aureus* strain Newman (Newman SmR) at 2×10^8^ colony forming units (CFU)/nostril, or PBS. Over the course of 49 days, NT was harvested from mice, and absolute numbers of IL-17+ CD4+ T-cells present within the NT were assessed. There was an increase in the numbers of IL-17+ CD4+ T-cells at 7 days post-*S. aureus* exposure (Figure 1A and B), agreeing with previous studies that have documented an acute IL-17 T-cell response in the NT following a single *S. aureus* exposure [11]. This IL-17 CD4+ T-cell response was reduced prior to complete clearance of *S. aureus* from the NT, by day 42 post-colonisation (Figure 1C). To investigate if T-cells recruited to the NT during *S. aureus* colonisation gave rise to a resident memory T-cell population, wild-type C57BL6/JCrl mice that were i.n. inoculated with *S. aureus* strain Newman SmR, received a secondary i.n. administration of either *S. aureus*, on day 56 post-initial exposure. CD4+ T-cell expansion in the NT of 56 day “previously exposed” mice was assessed 3 days post-secondary *S. aureus* exposure (Figure 1D). To determine tissue residency, a previously defined intravenous (i.v) labelling approach was used [26], where mice were injected i.v. with a fluorochrome-labelled anti-CD45 antibody 10 minutes prior to euthanasia to stain circulating lymphocytes and allow for negative selection of tissue resident cells (Figure 1D). NT samples were stained for tissue residency markers CD44 and CD69 (Supplemental 1). Proportions of total live cells, total CD4+ T-cells, CD4+ T-cells expressing surface markers of tissue resident memory (CD44high, CD69+), and IL-17+ T-cells expressing tissue residency memory markers (CD4+, CD44high, CD69+) within the NT that were negative for IVCD45 were assessed in these mice (Figure 1E and F). While total live cells and CD4+ T-cells in the NT had some cells that were positive for the i.v. injected CD45, the proportion of CD4+ TRM cells (CD44high, CD69+) and IL-17+ CD4+TRM cells that were IVCD45-was close to 100% in all samples (Figure 1E), indicating tissue residency. As such, expression of markers CD69 and CD44 was deemed appropriate to denote tissue residency in NT samples in future experiments.

**Figure 1.**
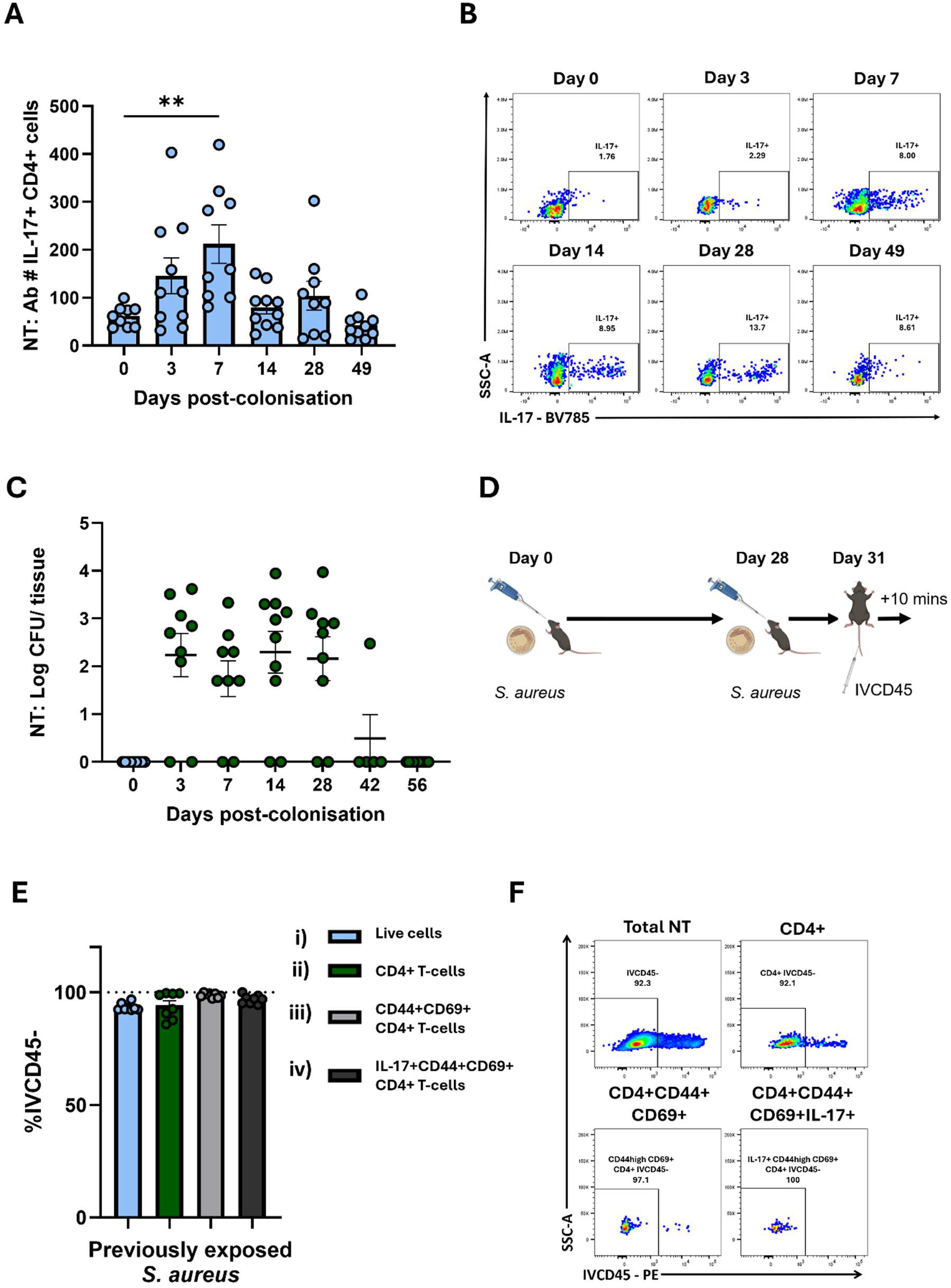
*S. aureus* colonisation drives expansion of IL-17+ CD4+T cells in the NT. Wild-type C57BL6/JCrl SPF mice were i.n. inoculated with *S. aureus* Newman SmR (2×10^8^ CFU/nostril in 10µl) or PBS on day 0. At specific time points NT was harvested and tissue digested for flow cytometry analysis. Cells were gated on CD45+, CD3+, CD4+cells. The absolute cell number (Ab#) of IL-17 producing CD4+ T-cells were assessed at days 3, 7, 14, 28 and 49 post-colonisation (A). Representative flow plots of IL-17+ CD4+ T-cells are shown (B). NT was homogenized and serial dilutions of homogenates were plated onto streptomycin-supplemented TSA plates. Plates were grown overnight and *S. aureus* CFU were enumerated on days 3, 7, 14, 28, 49 and 52 post-colonisation. Results are expressed as Log CFU/tissue (C). Previously exposed mice were re-exposed to *S. aureus* on day 28 post-initial *S. aureus* exposure. 3 days after secondary *S. aureus* exposure, these previously exposed mice were i.v. administered a CD45 fluorochrome-labelled antibody, and 10 minutes later NT was removed (D). The percentage (%) of total NT-cells (i), CD4+ cells (ii), CD4+ CD44high CD69+ cells (iii) and IL-17+ CD4+ CD44high CD69+ cells (iv), that are IVCD45-in the NT were assessed (E). Representative flow plots of IVCD45 expression by total NT cells, CD4+ cells, CD4+ CD44high CD69+ cells and IL-17+ CD4+ CD44high CD69+ cells are shown (F). Results are expressed as mean ± S.E.M. (n=5-10 per group). Statistical analysis was performed using one-way analysis of variance. **P ≤ 0.01

To assess re-activation of resident CD4+ T-cells in the NT, “naive” mice that received PBS, and “previously exposed” mice that received *S. aureus* 56 days prior were exposed to either PBS or *S. aureus* (Figure 2A). Re-exposure of mice to *S. aureus* resulted in a significant increase in the numbers of NT CD4+ TRM (CD44high, CD69+) (Figure 2B and D), and CD4+ TRM cells producing IL-17 (Figure 2C and E) on day 3 post re-exposure. CD4+ TRM cells in previously exposed mice also displayed significantly higher Ki67 expression on day 3 post re-exposure compared to naive mice that were administered *S. aureus* and previously exposed mice that received PBS (Supplemental 2A and B). This suggests that while these cells retract to baseline after *S. aureus* clearance, the increase in CD4+ TRM cells upon re-exposure to *S. aureus* is likely due to in situ expansion. To further establish that CD69+ CD44high CD4+ T cells that are re-activated upon secondary *S. aureus* exposure are not infiltrating lymphocytes, one group of previously exposed mice was administered FTY720. This sphingosine-1-phosphate receptor 1 (S1PR1) agonist that inhibits lymphocyte egress from lymph nodes (LNs) [27] was administered in drinking water 10 days prior to secondary *S. aureus* exposure and for the duration of the experiment (Figure 2A). When NT were assessed on day 3 post-re-exposure, there was no difference in the numbers of total CD4+ TRM (CD44high, CD69+) (Figure 2F and H) or IL-17+ CD4+ TRM (Figure 2G and I) cells in the previously exposed mice that received FTY720 prior to secondary *S. aureus* exposure compared to mice that did not receive FTY720. These findings indicate that the increase in cell number upon re-exposure to *S. aureus* is not dependent on lymphocyte infiltration, and CD44high CD69+ CD4+ that are re-activated upon re-exposure to *S. aureus* are locally proliferating TRM cells. As an internal control to assess if FTY720 successfully inhibited lymphocyte infiltration, proportions of CD44-CD4+ T-cells in the NT were compared, as naïve CD44-T-cells are known to be depleted upon successful administration of FTY720 [28]. Proportions of NT CD4+ T-cells that were CD44-were significantly lower in previously exposed mice that received FTY720 prior to secondary *S. aureus* exposure compared to those that did not (Supplemental 2C and D), indicating that FTY720 administration had successfully depleted circulating lymphocyte populations [28]. Re-exposure of previously exposed mice to *S. aureus* also resulted in a significant increase in the numbers of total (Figure 2J and L), and IL-17+ (Figure 2K and M) CD44high, CD27-γδ+ T-cells in the NT 3 days post re-exposure. This suggests that *S. aureus* colonisation similarly expands a population of memory γδ+ T-cells in the NT.

**Figure 2.**
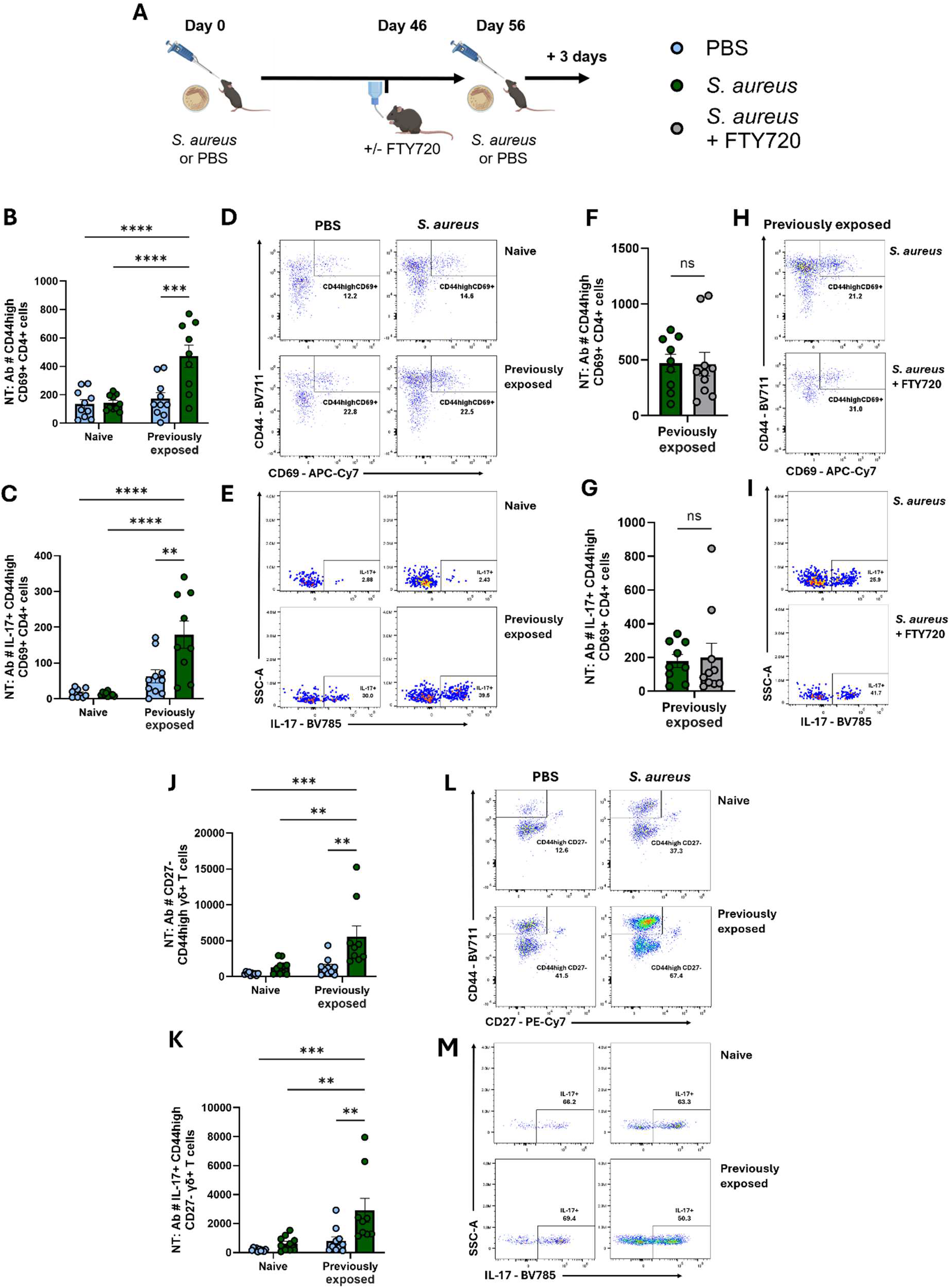
*S. aureus* colonisation drives expansion of IL-17+ CD4+TRM cells and γδ+ TRM cells in the NT. Wild-type C57BL6/JCrl SPF mice were i.n. inoculated with *S. aureus* Newman SmR (2×10^8^ CFU/nostril in 10µl) or PBS on day 0. Previously exposed or naive mice were re-exposed to *S. aureus* on day 56 post-initial *S. aureus* exposure. One group of previously exposed mice received FTY720 in drinking water 10 days prior to secondary *S. aureus* administration and for the duration of the experiment. 3 days after secondary *S. aureus* exposure NT was harvested (A). The absolute cell numbers (Ab #) of CD4+ TRM cells (CD4+, CD44high, CD69+) (B), and IL-17+ CD4+ TRM (IL-17+ CD4+ CD44high CD69+) (C), were assessed in the NT of naive and previously exposed mice following re-exposure to *S. aureus* or PBS. Representative flow plots of CD4+ TRM cells (D) and IL-17+ CD4+ TRM cells are shown (E). The absolute cell number (Ab #) of CD4+ TRM cells (F), and IL-17+CD4+ TRM cells (G) in the NT of naive and previously exposed mice that received FTY720 prior to *S. aureus* re-exposure were also assessed and compared to the previously exposed mice that did not receive FTY720. Representative flow plots of CD4+ TRM cells (H) and IL-17+CD4+ NT TRM cells are shown (I). The absolute cell numbers (Ab #) of γδ+ TRM cells (γδ+, CD44high, CD27-) (J), and IL-17+ γδ+ TRM cells (IL-17+ γδ+, CD44high, CD27-) (K), were assessed in the NT of naive and previously exposed mice following re-exposure to *S. aureus* or PBS. Representative flow plots of γδ+ TRM cells (L) and IL-17+ γδ+ TRM cells are shown (M). Results are expressed as mean ± S.E.M. (n=9-10 per group). Statistical analysis was performed using two-way analysis of variance or student t test. **P ≤ 0.01, ***p ≤ 0.001, ****p ≤ 0.0001.

Together, these findings suggest that both CD4+ and γδ+ TRM cells remain primed in the NT following *S. aureus* colonisation in the absence of the bacterium, with the potential for local expansion and increased cytokine production upon secondary *S. aureus* exposure. Notably re-activation of IL-17 producing TRM cells coincided with enhanced clearance of *S. aureus* from the Nose (Supplemental 2E) and NT (Supplemental 2F) in previously exposed mice, 3 days following re-exposure.

### *S. aureus* colonisation drives CD4+ TRM cell expansion in the NT in the absence of other microbes

Unlike humans that are commonly persistently colonised with *S. aureus* in the anterior nares, mice that receive a bolus i.n. administration of *S. aureus* have previously been shown to clear the bacteria from the NT approximately 28-42 days post-colonisation [13],[29]. To establish a model of long-term persistent colonisation, *S. aureus* Newman was introduced into the bedding of germ-free (GF) mice (Figure 3A). This environmental exposure resulted in mice that remained nasally colonised with *S. aureus* up to 28 days post-mono-colonisation (Figure 3B). Importantly, 6–12-week first generation (F1) progeny of these mice were stably colonised with *S. aureus* in the nose (Figure 3B) NT and colon (Figure 3C) with limited detection of *S. aureus* in other tissues (Figure 3C). Mono-colonisation of GF mice with *S. aureus* resulted in a significant increase in the numbers of CD4+ T-cells expressing TRM markers (CD44high, CD69+) within the NT in 6–12-week-old F1 *S. aureus* mono-colonised mice compared to GF mice (Figure 3D and E). Levels of CD4+ TRM cells producing IL-17 in the NT of F1 mono-colonised mice were also significantly higher than in GF mice (Figure 3F and G), similar to what was seen in previously exposed conventional specific pathogen free (SPF) mice 2 months post transient colonisation (Figure 3F and G). This indicates that both direct nasal colonisation of SPF mice and mono-colonisation of GF mice through environmental exposure to *S. aureus,* expands IL-17+ CD4+ TRM cells in the NT. Interestingly, there was no increase in the proportions of IL-17+, CD4+ TRM in the spleen or LNs (Figure 3H and I) in *S. aureus* mono-colonised mice compared to GF mice, suggesting that the presence of *S. aureus* alone drives site specific CD4+ TRM cell expansion.

**Figure 3.**
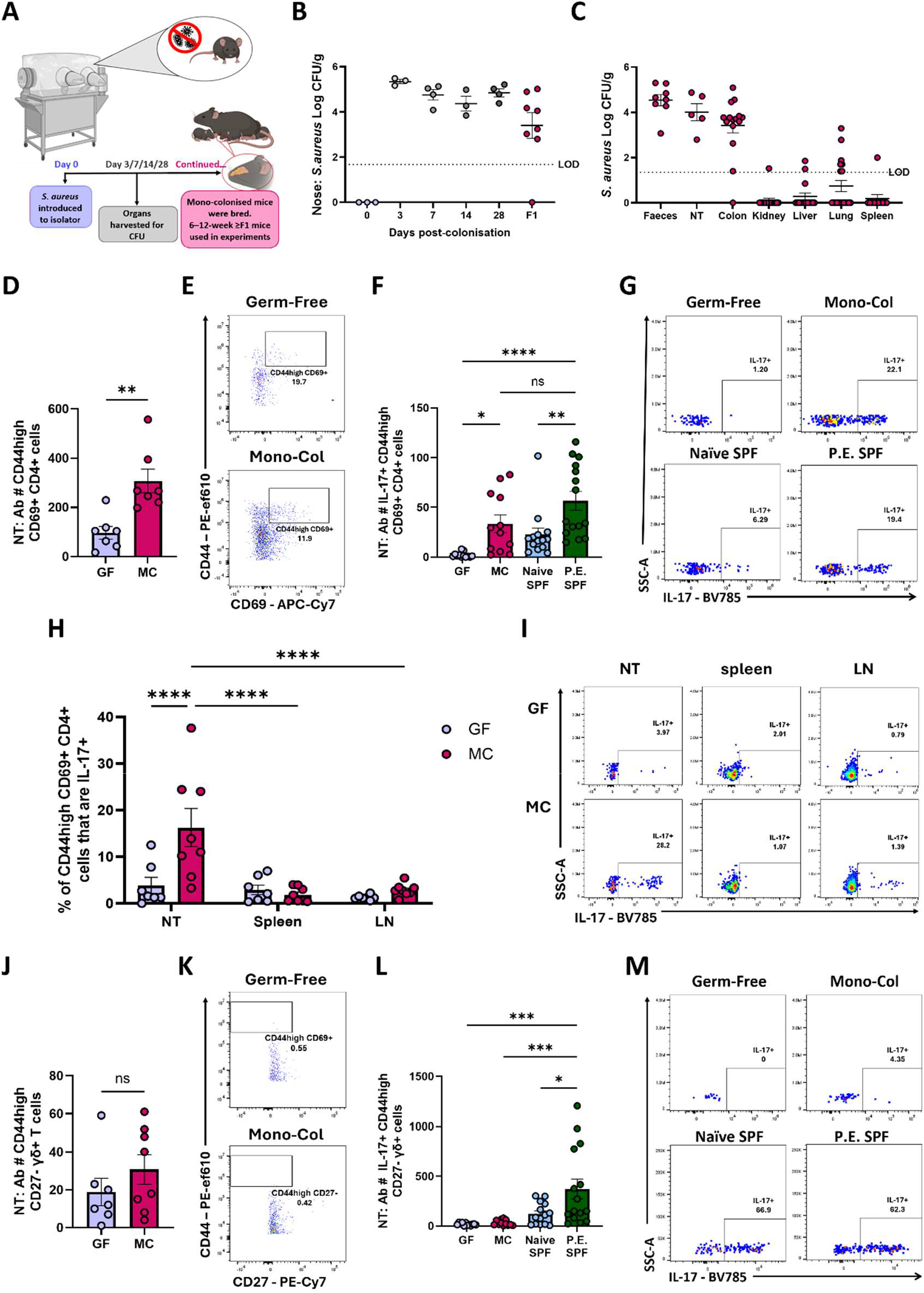
*S. aureus* mono-colonisation of Germ-Free mice expands CD4+, but not γδ+,TRM cells in the NT. *S. aureus* strain Newman was introduced into the bedding that housed GF mice to generate *S. aureus* mono-colonised mice that were persistently colonised (A). Nose tissue was harvested from GF mice at days 0, 3, 7, 14, and 28 post-introduction of *S. aureus* into the environment, as well as from first generation (F1) mono-colonised mice, and *S. aureus* bacterial burden, expressed as log CFU/g of tissue, assessed (B). Organs were also harvested from F1 *S. aureus* mono-colonised mice and *S. aureus* bacterial burden, expressed as log CFU/g of tissue, assessed in colon, NT, lung, spleen, liver, and kidney as well as in the faeces (C). NT, spleen and mediastinal lymph nodes (LN) were harvested from 6–12-week (≥F1 generation) GF and *S. aureus* mono-colonised mice and tissue was digested for flow cytometry analysis. Cells were gated on CD45+, CD3+ cells. The absolute cell number (Ab #) of CD4+ TRM cells (CD4+, CD44high, CD69+) in the NT of 6–12-week GF and F1 *S. aureus* mono-colonised mice were assessed (D). Representative flow plots of NT CD4+ TRM cells are shown (E). The absolute cell number (Ab #) of IL-17+ CD4+ TRM cells (IL-17+, CD4+, CD44high, CD69+) in the NT of GF, 6-12 weeks (F1 generation) *S. aureus* mono-colonised mice was compared to naïve SPF, mice and previously exposed SPF mice, 28-day post *S. aureus* colonisation (P.E.) (F). Representative flow plots of NT IL-17+ CD4+ TRM cells are shown (G). Percentage (%) of IL-17+ CD4+ TRM cells in the NT, spleen, and LNs, of GF and 6–12-week F1 *S. aureus* mono-colonised mice were assessed (H). Representative flow plots of IL-17+CD4+ TRM cells from NT, spleens and LNs of GF and 6–12-week F1 *S. aureus* mono-colonised mice are shown (I). The absolute cell number (Ab #) of γδ+ TRM cells (γδ+, CD44high, CD27-) in the NT of GF and 6–12-week F1 *S. aureus* mono-colonised mice were assessed (J). Representative flow plots of NT γδ+ TRM cells are shown (K). The absolute cell number (Ab #) of IL-17+ γδ+ TRM cells (IL-17+, γδ+, CD44high, CD27-) in the NT of GF, F1 *S. aureus* mono-colonised, naïve SPF, and 28-day previously exposed (P.E.) SPF mice were assessed (L). Representative flow plots of NT L-17+ γδ+ TRM cells are shown (M). Results are expressed as mean ± S.E.M. (n=3-19 per group for CFU, n=3-8 for flow cytometry). Statistical analysis was performed using one-way or two-way analysis of variance, or student t test. *P ≤ 0.05, **p ≤ 0.01, ***p ≤ 0.001, ****p ≤ 0.0001.

Conversely, CD44high, CD27-γδ+ T-cells existed at relatively low numbers in both 6–12-week-old F1 mono-colonised mice and GF controls, with no significant difference in cell numbers between groups (Figure 3J and K). When compared to SPF mice that had been previously exposed to *S. aureus,* both mono-colonised and GF mice had significantly lower numbers of IL-17+ CD44high, CD27-γδ+ T-cells (Figure 3L and M), suggesting that mono-colonisation with *S. aureus* is not sufficient to maintain a population of γδ+ T-cells in the NT.

This data indicates that the presence of *S. aureus* in the absence of a complex microbiome is sufficient to drives site specific expansion of CD4+, but not γδ+, NT resident T-cells.

### *S. aureus* colonisation expanded NT CD4+ TRM cells provide protection during heterologous infection

Having demonstrated that nasal colonisation with *S. aureus* drives NT CD4+ TRM cell expansion, the capacity of these CD4+ TRM cells for non-specific reactivation was assessed. Wild-type C57BL6/JCrl SPF mice were i.n. inoculated with *S. aureus* strain Newman SmR or PBS, then left for 28 days. Naive mice that had received PBS 28 days prior, and previously exposed mice that had received *S. aureus* 28 days prior, were i.n. administered LPS at 10µg/nostril in 10µl, or PBS, and left for a further 4- or 72-hours (Figure 4A). There was a rapid and significant increase in the number of CD4+ TRM cells (Figure 4B and C) and IL-17+ CD4+ TRM cells (Figure 4D and E) in the NT of mice previously exposed to *S. aureus* mice that received LPS, compared to naive mice that received LPS or previously exposed mice that received PBS. Rapid re-activation of these cells as early as 4 hours post LPS stimulation supports the theory that *S. aureus* conditioned-CD4 TRM cells are sitting primed in the NT to rapidly respond at this exposed barrier site.

**Figure 4.**
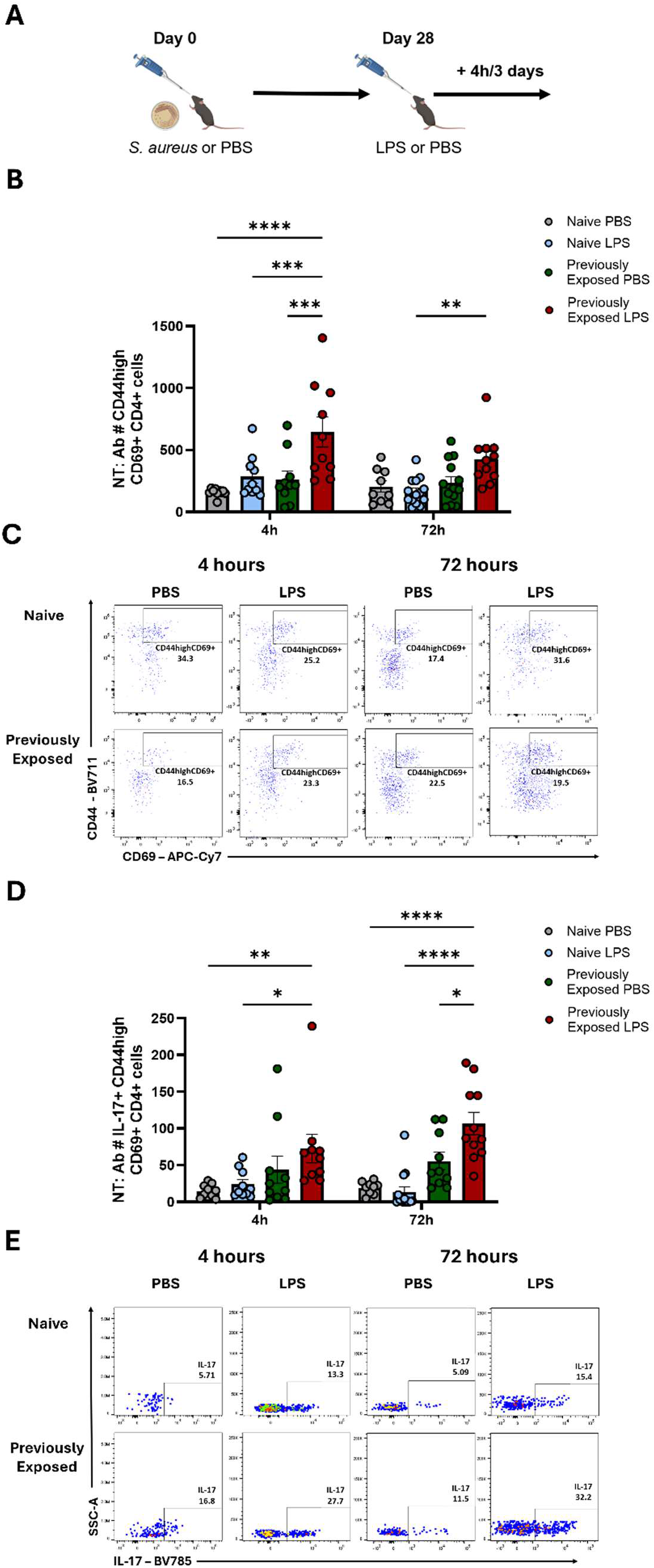
NT IL-17+ CD4+ TRM cells in *S. aureus* colonised mice can be activated non-specifically by LPS. Wild-type C57BL6/JCrl SPF mice were i.n. inoculated with *S. aureus* (Newman SmR 2×10^8^ CFU/nostril in 10µl) or PBS. LPS or PBS was administered i.n. to previously exposed mice (28-days post-*S. aureus* exposure), or to naive control mice. At specific timepoints post-secondary i.n. challenge, NT were harvested for flow cytometric analysis. The absolute cell number (Ab #) of CD4+ TRM (CD4+, CD44high, CD69+) (B) and IL-17+ CD4+ TRM (D) cells in the NT of 28-day previously exposed or naive mice at 4- and 72-hours post-challenge with LPS or PBS were assessed. Representative flow plots of NT CD4+ TRM cells (C) and IL-17+ CD4+ TRM (E) are shown. Results are expressed as mean ± S.E.M. (n=8-14 per group). Statistical analysis was performed using one-way or two-way analysis of variance. *P ≤ 0.05, **p ≤ 0.01, ***p ≤ 0.001, ****p ≤ 0.0001.

To investigate the significance of this non-specific CD4+ TRM cell reactivation in the context of infection, wild-type C57BL6/JCrl SPF mice were i.n. inoculated with *S. aureus* strain Newman SmR or PBS, then left for 56 days. Naive mice that had received PBS 56 days prior, and previously exposed mice that had received *S. aureus* 56 days prior, were then i.n. challenged with *K. pneumoniae*, or PBS, and left for a further 24- or 72-hours (Figure 5A). CD4+ TRM cell (Figure 5B and D) and IL-17+ CD4+ TRM cell (Fig 5C and D) numbers increased in the NT of previously exposed mice infected with *K. pneumoniae* compared to naive, *K. pneumoniae* infected mice, at 72-hours post-infection. This increase in CD4+ TRM cell number and IL-17 production was accompanied by a significant reduction in bacterial burden in the NT (Figure 5E) and nose (Figure 5F) of *K. pneumoniae* challenged previously exposed mice compared to the *K. pneumoniae-*challenged naive mice, at 72-hours post-infection. One group of previously exposed mice also received FTY720 in their drinking water 10 days prior to the *K. pneumoniae* challenge (Figure 5A). Additionally, in one group of previously exposed mice IL-17 production was inhibited by i.n. administration of anti-IL-17 mono-clonal antibody alongside *K. pneumoniae* infection, as well as a secondary administration of anti-IL-17 antibody 24-hours post-infection. *K. pneumoniae* clearance from the NT (Figure 5E) and lung (Figure 5G) was impaired in previously exposed mice by IL-17 blocking, but in the NT (Figure 5E) and nose (Figure 5F) *K. pneumoniae* clearance was not affected by FTY720 treatment. These findings suggest that the enhanced clearance of *K. pneumoniae* observed in *S. aureus* previously exposed mice is dependent on IL-17, but independent of lymphocyte infiltration. IL-17 has been shown to enhance *K. pneumoniae* clearance by driving the recruitment of neutrophils to infected tissue [30]. Lungs were therefore analysed by flow cytometry 3 days post-*K. pneumoniae* infection, gating on CD45+, Ly6G+, CD11b+ cells (Supplemental 3). Trends indicated that both numbers and proportions of neutrophils recruited to the lung begin to decrease by 24 hours post-infection (Figure 5H-J) and are significantly reduced by 72 hours post-infection (Figure 5K-M) when IL-17 was blocked in previously exposed, *K. pneumoniae* infected mice. This suggests that the enhanced protection observed in the *S. aureus* previously exposed mice could be due to IL-17+ TRM cells promoting neutrophil recruitment to the tissue.

**Figure 5.**
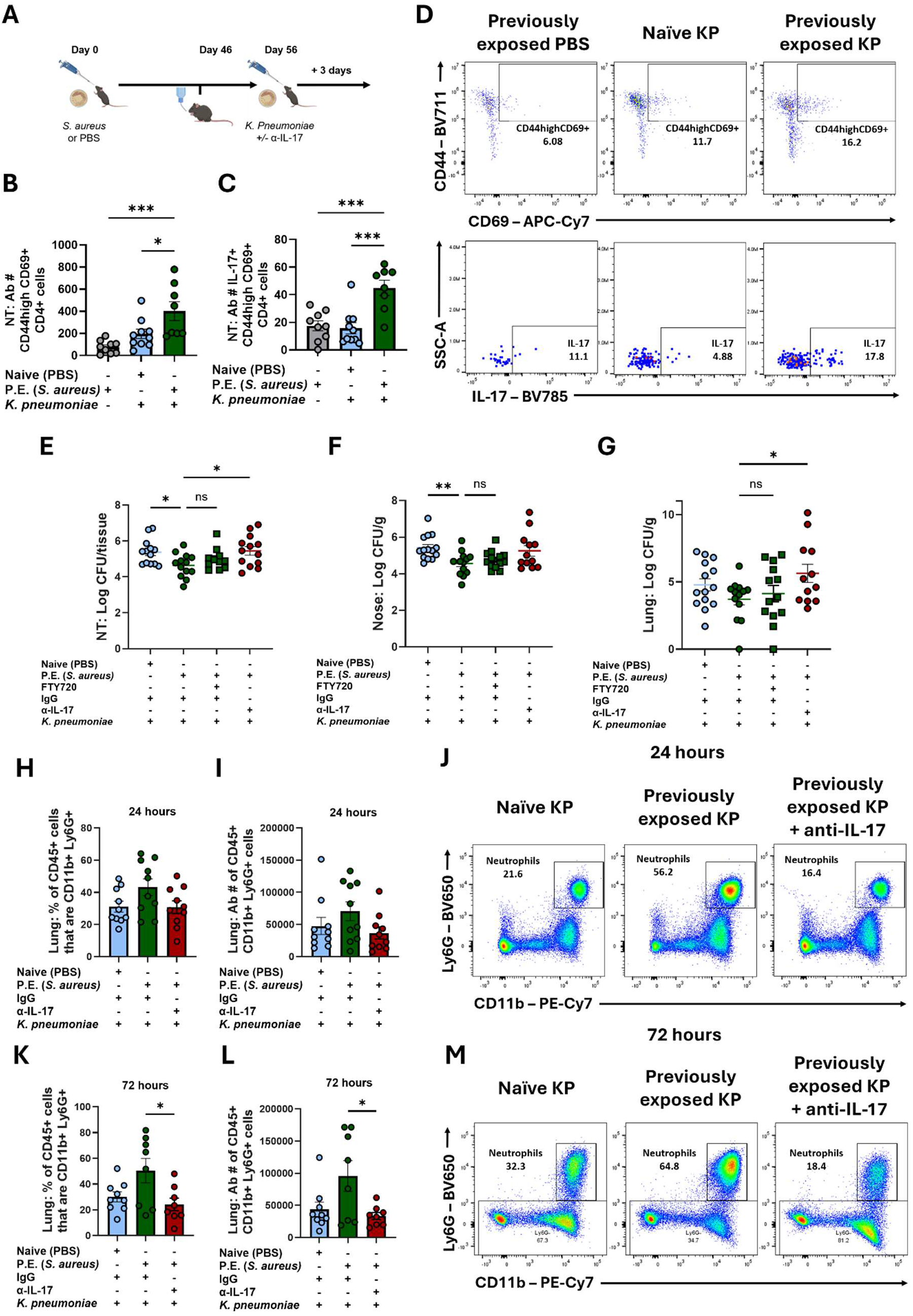
NT IL-17+ CD4+ TRM cells generated in *S. aureus* colonised mice can be activated non-specifically during *K. pneumoniae* infection. Wild-type C57BL6/JCrl mice were i.n. inoculated with *S. aureus* (Newman SmR 2×10^8^ CFU/nostril in 10µl) or PBS. PBS or *K. pneumoniae* (Strain ATCC Kp43816 2.5X10^3^CFU/ nostril in 25μl) was administered i.n. to previously exposed mice (56-days post-*S. aureus* exposure), or to naive control mice. Mice were administered either an anti-IL-17 mono-clonal antibody (α-IL-17) or an isotype control (IgG) alongside i.n. *K. pneumoniae* inoculum, as well as a secondary administration 24 hours later. One group of previously exposed mice received FTY720 in drinking water 10 days prior to *K. pneumoniae* administration and for the duration of the experiment. 24 and 72 hours post-secondary i.n. challenge, NT and lung were harvested for flow cytometric analysis (A). The absolute cell number (Ab #) of CD4+ TRM (CD4+, CD44high, CD69+) (B) and IL-17+ CD4+ TRM (C) cells in the NT of 56-day previously exposed or naive mice, on day 3 post-infection with *K. pneumoniae* or PBS was assessed. Representative flow plots of NT CD4+ TRM cells and IL-17+ CD4+ TRM are shown are shown (D). *K. pneumoniae* bacterial burden expressed as log CFU per total tissue sample in the NT (E), or as log CFU/g of tissue in the nose (F) and Lung (G), was assessed in naive or 56-day previously exposed mice +/− FTY720 that received anti-IL-17 or an isotype control, 3 days post-infection with *K. pneumoniae*. The percentage (H and K) and absolute cell number (Ab #) (I and L) of neutrophils (CD45+, CD11b+, Ly6G+) in the lung of naive or 56-day previously exposed mice +/− anti-IL-17 administration at 24- (H and I) or 72- (K and L) hours post-infection with *K. pneumoniae* was assessed. Representative flow plots of neutrophils in the lung 24- (J) or 72- (M) hours post-infection with *K. pneumoniae* are shown. Results are expressed as mean ± S.E.M. (n=8-14 per group). Statistical analysis was performed using one-way or two-way analysis of variance. *P ≤ 0.05, **p ≤ 0.01, ***p ≤ 0.001.

To investigate if CD4+ TRM cells generated in response to *S. aureus* colonisation can provide heterologous protection in the absence of γδ+ T-cells, 6–12-week F1 S. aureus mono-colonised mice, along with GF controls, were i.n. challenged with *K. pneumoniae* (Figure 6A). Mice were weighed daily for 3 days post-infection. GF mice lost significantly more weight post-*K. pneumoniae* infection compared to *S. aureus* mono-colonised mice (Figure 6B). At 24- and 72-hours post-infection, organs were harvested for CFU enumeration. *S. aureus* mono-colonised mice had significantly lower *K. pneumoniae* bacterial burden in the lungs at both 24- and 72-hours post-infection (Figure 6C) and significantly lower bacterial burden in the NT (Figure 6D), nose (Figure 6E) and liver (Figure 6F) at 72-hours post-infection compared to GF controls (Figure 6D-F). *K. pneumoniae* bacterial burden was also reduced in the kidneys of *S. aureus* mono-colonised mice compared to GF controls, but not significantly (Figure 6G). This increased clearance of *K. pneumoniae* was associated with a significant increase in the number of IL-17+, CD4+ TRM cells in the *S. aureus* mono-colonised NT compared to GF controls at 72-hours post-*K. pneumoniae* challenge (Figure 6H and I). Importantly, there was no significant difference in the number of IL-17+ γδ+ T-cells between GF and *S. aureus* mono-colonised mice (Figure 6J and K). I.n. administration of an anti-IL-17 mono-clonal antibody alongside *K. pneumoniae* infection, and at 24-hours post-infection, in *S. aureus* mono-colonised mice (Figure 6L), resulted in significant impairment of *K. pneumoniae* clearance in the lung (Figure 6M), NT (Figure 6N), nose (Figure 6O) and liver (Figure 6P) 72-hours post-infection.

**Figure 6.**
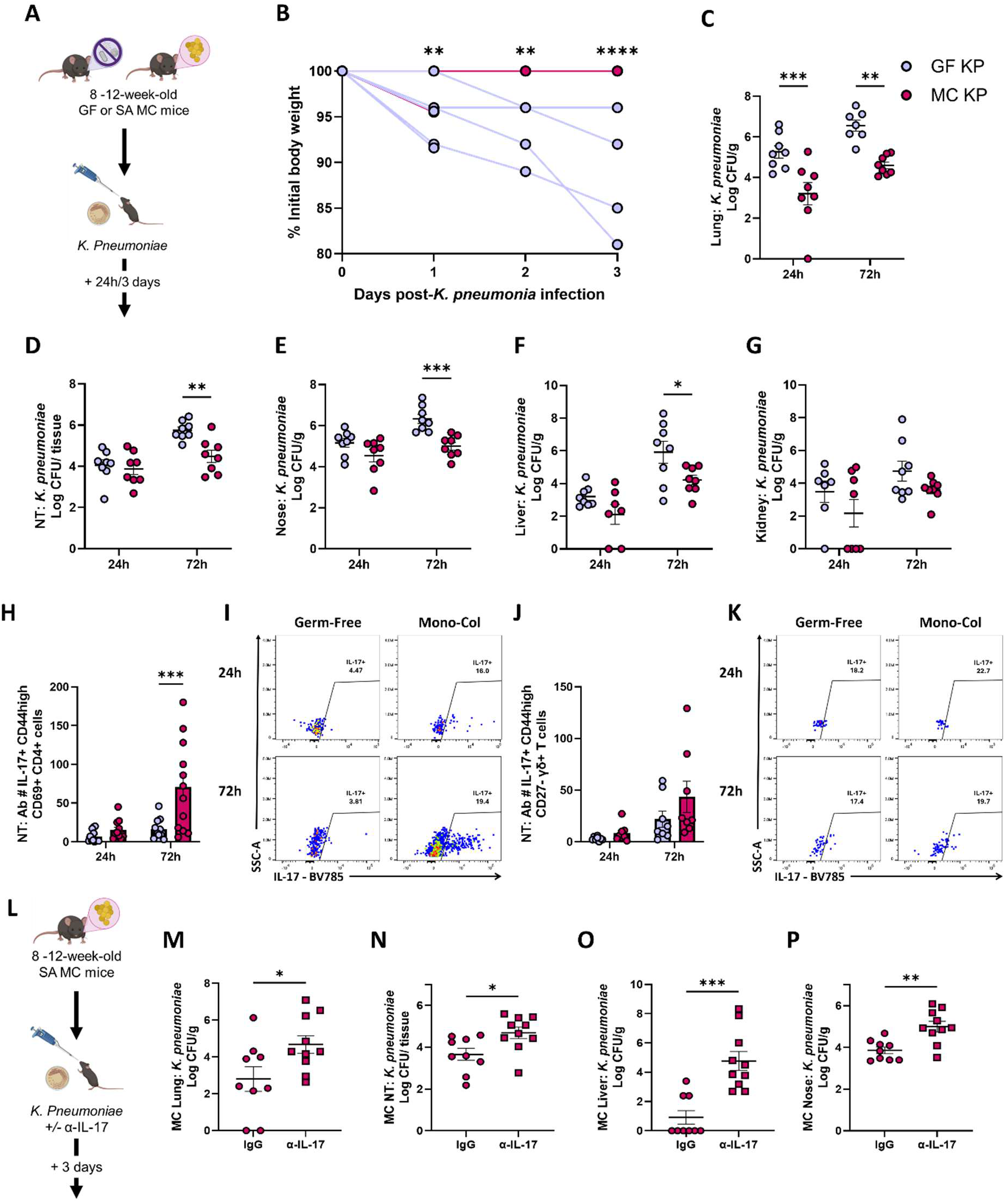
*S. aureus* mono-colonisation confers protection against *K. pneumoniae* infection. *S. aureus* strain Newman was introduced into the bedding that housed GF mice to generate a population of *S. aureus* mono-colonised mice that were persistently colonised from birth. *K. pneumoniae* (Strain ATCC Kp43816 2.5X10^3^CFU/ nostril in 25μl) or PBS, was administered i.n. to 6–12-week *S. aureus* mono-colonised and GF mice, at least F1 generation (A). Body weight was assessed and expressed as a percentage of pre-infection weight at 24- and 72-hours post-*K. pneumoniae* infection (B). 24 and 72 hours post-*K. pneumoniae* infection tissues were harvested and processed for CFU enumeration. *K. pneumoniae* bacterial burden expressed as log CFU/g of tissue in the lung (C), nose (E), liver (F) and kidney (G), or CFU/g total tissue in the NT (D), was assessed 24- and 72-hours post-infection with *K. pneumoniae*. NT were digested for flow cytometry analysis. Cells were gated on CD45+, CD3+ cells. The absolute cell number (Ab #) of IL-17+ CD4+ TRM cells (H) and IL-17+ γδ+ TRM cells (J) in the NT of GF and *S. aureus* mono-colonised mice were assessed 24- and 72-hours post-infection with *K. pneumoniae.* Representative flow plots of NT IL-17+ CD4+ TRM cells (I) IL-17+ γδ+ TRM cells (K) and are shown. In a second experimental setup, *K. pneumoniae* (Strain ATCC Kp43816 2.5X10^3^CFU/ nostril in 25μl) or PBS, was administered i.n. to 6–12-week *S. aureus* mono-colonised and GF mice, at least F1 generation. *S. aureus* mono-colonised mice were administered either an anti-IL-17 mono-clonal antibody (αIL-17) or an isotype control (IgG) alongside i.n. *K. pneumoniae* challenge, as well as a secondary administration 24 hours later (L). *K. pneumoniae* bacterial burden expressed as log CFU/g of tissue in the lung (M), nose (O), and liver (P), or CFU/g total tissue in the NT (N), was assessed in *S. aureus* mono-colonised mice that were administered anti-IL-17 or an isotype control, 72-hours post-infection with *K. pneumoniae*. Results are expressed as mean ± S.E.M. (n=4-12) Statistical analysis was performed using two-way analysis of variance or student t test. *P ≤ 0.05, **p ≤ 0.01, ***p ≤ 0.001, ****p ≤ 0.0001.

Overall, these findings demonstrate that nasal colonisation with *S. aureus* enhances protection during heterologous *K. pneumoniae* infection and suggest that IL-17 production by NT CD4+ TRM cells play a critical role in this protection.

### *S. aureus* expanded CD4+ TRM cells in the NT can be bystander activated

To investigate how NT CD4+ TRM cells from *S. aureus* previously exposed mice are reactivated during *K. pneumoniae* infection, NT were harvested from C57BL6/JCrl SPF mice 28 days post i.n inoculation with *S. aureus* strain Newman SmR or PBS, and a number of ex-vivo investigations were performed (Supplemental 4). Initially, total NT cells from previously exposed and naive mice were cultured ex-vivo for 72-hours in the presence of heat-killed *S. aureus* or heat-killed *K. pneumoniae*. Both stimulations resulted in significantly more IL-17 production from previously exposed compared to naive mouse NT cells (Figure 7A). This increased cell reactivation was specific to NT cells and not observed when splenocytes from previously exposed mice were challenged with heat-killed *S. aureus* or *K. pneumoniae* (Figure 7B). CD4+ T-cells were then isolated from the NT of *S. aureus* previously exposed or naive mice and cultured alongside irradiated splenic antigen presenting cells (APCs) for 72-hours. CD4+ T-cells from previously exposed mice responded to stimulation with heat-killed *S. aureus,* but not heat-killed *K. pneumoniae,* (Figure 7C), suggesting that CD4+ T-cells from the NT of previously exposed mice are *S. aureus* specific, and *K. pneumoniae* antigen presentation is not sufficient to reactivate these cells. Previous studies have shown that bystander activation of CD4+ NT TRM cells in other infection models is mediated by cytokines, likely produced by innate immune cells that are activated during heterologous infection [16]. Consistent with this, CD4+ TRM cells isolated from *S. aureus* previously exposed mice could be activated in vitro by Th17 polarising cytokines in the absence of antigen (Figure 7D). Treatment with IL-23 and IL-7 in conjunction with either IL-1β or IL-18 resulted in significantly more IL-17 production by CD4+ TRM cells from previously exposed mice compared to CD4+ TRM from naive mice (Figure 7D). Furthermore, when cell free supernatants from *K. pneumoniae-*treated bone marrow-derived dendritic cells (BMDCs) were used to stimulate NT CD4+ T-cells purified from naive and *S. aureus* previously exposed mice, NT CD4+ T-cells from *S. aureus* previously exposed mice produced significantly more IL-17 (Figure 7E). BMDCs were confirmed to produce IL-1β and IL-23 following exposure to live *K. pneumoniae,* but not IL-12 (Figure 7F). Finally, it was shown that IL-17 production by NT CD4+ T-cells from *S. aureus* previously exposed mice stimulated with *K. pneumoniae-*treated BMDC cell free supernatants could be significantly decreased when IL-23 was blocked and further reduced when IL-23 and IL-1β were blocked in unison (Figure 7G).

**Figure 7.**
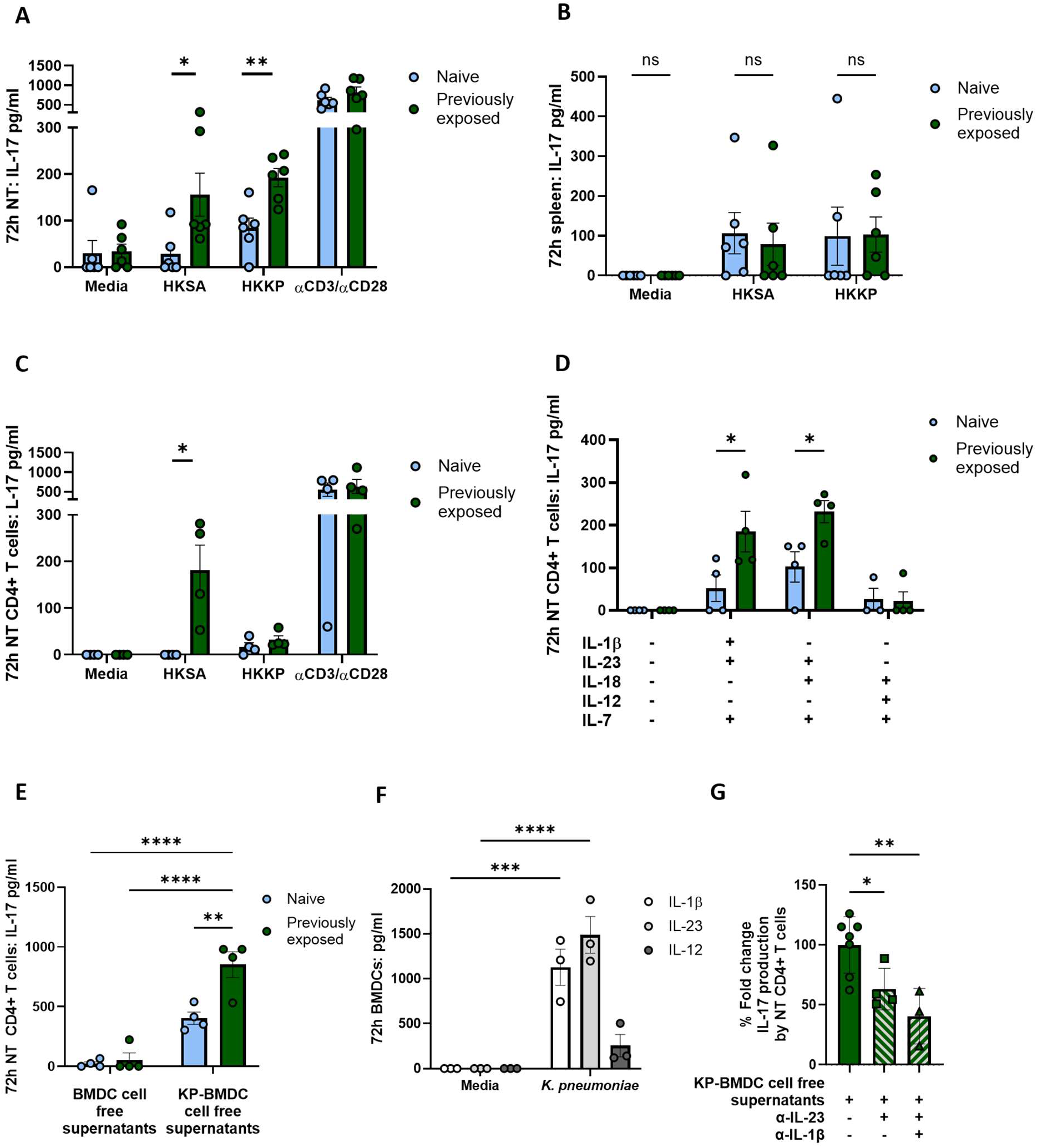
NT CD4+ T-cells generated in *S. aureus* colonised mice can be bystander activated. Wild-type C57BL6/JCrl mice were intranasally inoculated with *S. aureus* (Newman SmR 2×10^8^ CFU/nostril in 10µl) or PBS. At day 28 post-colonisation, NT was harvested and digested. Total NT-cells (1×10^5^ cells/well) (A) or total splenocytes (1×10^5^ cells/well) (B) were stimulated in vitro with heat-killed *S. aureus* or heat killed *K. pneumoniae* (1μg/ml) for 3 days before IL-17 secretion was assessed by ELISA and expressed as pg/ml. NT samples isolated from Naive and 28-day previously exposed mice were MACs sorted for CD4+ T-cells, which were then cultured in vitro (1×10^4^ cells/well) in the presence of irradiated splenocytes (1×10^4^ cells/well), which acted as antigen presenting cells, and were stimulated with heat-killed *S. aureus* or heat killed *K. pneumoniae*. IL-17 secretion was assessed by ELISA and expressed as pg/ml (C). NT CD4+ T-cells isolated from naive and 28-day previously exposed mice were cultured in the absence of APCs and stimulated with combinations of recombinant cytokines IL-1β, IL-23, IL-18, IL-12, IL-7 (10ng/ml) or media alone. IL-17 secretion was assessed on day 3 by ELISA and expressed as pg/ml (D). Murine bone marrow derived macrophages (BMDCs) were generated from naive mice and infected with *K. pneumoniae* MOI 50 for 1 hour, followed by gentamycin treatment for a further 23 hours. Cell free supernatant was collected and levels of IL-1β, IL-23 and IL-12 were assessed by ELISA (F). CD4+ T-cells isolated from naive and 28-day previously exposed NT samples, were cultured with irradiated splenocytes and stimulated with cell free supernatants collected from BMDCs infected with *K. pneumoniae* or media alone in the presence or absence of blocking antibodies against IL-23 or IL-23 and IL-1β. IL-17 secretion was assessed on day 3 by ELISA and expressed as pg/ml (G). Results are expressed as mean ± S.E.M. (n=8-12 mice per group, pooled from four independent experiments). Statistical analysis was performed using one-way or two-way analysis of variance. *P ≤ 0.05, **p ≤ 0.01, ***p ≤ 0.001, ****p ≤ 0.0001.

Overall, this data suggests that *S. aureus-*primed NT CD4+ TRM cells can undergo bystander activation, which is not achieved by antigen presentation and is reliant on cytokines produced by other immune cell populations at this tissue site.

## Discussion

Healthy individuals are frequently colonised with *S. aureus* in the anterior nares; however, the long-lasting immune consequences of asymptomatic colonisation are incompletely understood [6]. This study demonstrates that nasal colonisation with *S. aureus* primes IL-17 producing TRM cell populations in the NT. IL-17+ CD4+ TRM cells were expanded in the NT of both conventional and GF mice following *S. aureus* colonisation, while γδ+ TRM cells were only expanded in the NT of conventional *S. aureus* colonised mice. Intriguingly, TRM cells expanded in response to *S. aureus* had the capacity for non-specific reactivation and IL-17 production, which contributed to enhanced protection during subsequent infection with the respiratory pathogen *K. pneumoniae*. This enhanced protection was also observed in *S. aureus* mono-colonised mice that lack γδ+ T-cells, further supporting a critical role for CD4+ TRM cells in heterologous immune protection. Ex-vivo analysis demonstrated that heterologous reactivation of CD4+ TRM cells was due to TCR-independent bystander activation, driven by Th17 polarising cytokines, which could be produced by other local innate immune cell populations.

TRM cells are known to reside at mucosal sites acting as a first line of defence against invasive microbes [19]–[21]. While TRM cells have primarily been shown to be expanded during infection, studies in the skin [24] and gut [25] suggest that these cells can be shaped and maintained by commensal microbes that colonise these sites. This study now demonstrates that CD4+ and γδ+ TRM cell responses within the nasal mucosa are also shaped by bacterial colonisation. *S. aureus* colonisation of the murine NT drives upregulation of memory CD44high, CD69+, CD4+ T-cells, with close to 100% of these cells being negative for IVCD45 staining following *S. aureus* re-exposure. FTY720 administration prior to secondary *S. aureus* exposure further confirmed that the CD44high, CD69+, CD4+ memory T-cells were tissue resident and not circulating, supporting use of CD44 and CD69 expression to classify CD4+ TRM cells in the *S. aureus* colonised NT. While CD103 is also often expressed by TRM cells, these are primarily CD8+ TRM cells [31],[32]. As such, both CD103+ and CD103-TRM cells were included in all analysis. TRM cells in the respiratory tract are frequently reported to be potent IL-17 producers [33],[34], the primary cytokine reported to be produced during *S. aureus* colonisation [11],[12]. This study builds on our understanding of T-cell responses during *S. aureus* colonisation, indicating that, following initial T-cell activation, IL-17+ CD4+ and γδ+ TRM cells are retained in the tissue, acting as a first line of defence against other microorganisms that invade the nasal mucosa. A limitation of this flow cytometry data is our inability to confirm that these TRM cells are specific for *S. aureus* antigen. However, we do see higher levels of CD4+ T-cells re-activation when mice are re-exposed to *S. aureus* compared to naïve mice that are exposed to *S. aureus,* as well as *S. aureus* specific re-activation of these cells ex-vivo.

Findings here also demonstrate that *S. aureus* colonisation alone is not sufficient to retain γδ+ TRM cells in the NT of mono-colonised mice. This is in line with previous studies which show that γδ+ T cells require a diverse microbiome for persistence at mucosal surfaces and support the idea that specific signals derived from the microbiome shape the phenotype and function of local memory cell populations [35],[36]. Delineating specific bacterial associated signals that activate unique tissue resident memory cell populations is warranted.

NT CD4+ TRM cells expanded following colonisation with *S. aureus* have the capacity for non-specific expansion and IL-17 production upon subsequent exposure to *K. pneumoniae*, a key cytokine for neutrophil recruitment in *K. pneumoniae* clearance [37]. Findings here align with those of Curham et al., which demonstrated that NT CD4+ TRM cells have the capacity for bystander activation as early as 4 hours post-restimulation [16]. Ex vivo investigations demonstrated that NT CD4+ TRM cells isolated from *S. aureus* colonised mice could only be activated by *K. pneumoniae* antigen if other immune cell populations are present. Alternatively, these experienced TRM cells could be activated in the absence of TCR engagement if Th17 polarising cytokines were present, with optimal bystander activation achieved with IL-23 and IL-1β stimulation in conjunction with IL-7. IL-23 is known to be produced by dendritic cells following *K. pneumoniae* exposure [38], working synergistically with IL-1β to create a positive feedback loop which drives Th17-type cell activity [39]. Bystander activation of *S. aureus-*primed CD4+ TRM cells is likely not *K. pneumoniae* specific and may be observed during heterologous exposure to any microbe that induces Th17-polarising cytokine production locally within the tissue. It has previously been documented that antigen experienced TRM cells in peripheral barrier tissues exhibit heightened reactivation kinetics, which facilitates their activation in response to local inflammatory cytokine signals [39]. Findings here emphasise that *S. aureus* primed CD4+ TRM cells in the NT are ready for rapid re-activation to non-specific cytokine signalling which could clear other microbial populations from the *S. aureus* colonised tissue. A recent study from Aggarwal et al. suggests that persistent *S. aureus* colonisation is negatively associated with survival of three *Corynebacterium* species in the human nasal mucosa, suggesting that *S. aureus* has the capacity to outcompete other microbes to facilitate its persistence within this niche [40],[41]. NT TRM cells generated in response to colonising *S. aureus* that have the capacity for non-specific activation could be playing a critical role in this regard.

*S. aureus* has previously been shown to suppress host-immune responses to evade clearance in colonised individuals [8],[42]. Through the induction of IL-10 production by myeloid cells in the NT, *S. aureus* can create an anti-inflammatory environment that favours bacterial persistence [42]. Findings here, suggest that *S. aureus* can also drive immune stimulating effects during colonisation. The balance between these competing effects and how *S. aureus* could imprint immunological memory in both the innate and adaptive arms of the immune response is not understood and requires significant further investigation. However recent work suggests memory T cell plasticity may be a feature of prior *S. aureus* exposure [43]. To date, there has been no comprehensive report on whether *S. aureus* colonisation could be protective against infections with alternative bacterial species in humans. It is essential for these studies to be undertaken to fully appreciate the impact that *S. aureus* colonisation has on host immunity.

The complex interplay between *S. aureus* and the immune system means that *S. aureus* vaccine development remains a challenge [6]. This study emphasises the need to consider how any vaccines that fully eradicate *S. aureus* from the colonised host could impair TRM responses driven by colonisation. Additionally, harnessing the protective effects of pre-existing TRM cell populations in the respiratory tract of individuals with previous *S. aureus* exposure could represent an attractive approach to boost vaccine efficacy to protect against invasive *S. aureus* infection. For example, “prime and pull” vaccinations, which consist of a primary systemic vaccine followed by topical chemokine administration to “pull” memory T-cells to the infection site to induce localised protection [44] are thought to be more effective at sustaining long-term protection in immunised hosts [32]. This approach may not be directly applicable for *S. aureus* vaccination, as data here would suggest that protective TRM cell populations may already exist in the NT of individuals with previous *S. aureus* exposure. However, successful *S. aureus* immunisation may require dual administration of i.n. and systemic vaccines to support survival or further expansion of NT TRM cell populations but would require substantial additional investigation.

While the model used for *S. aureus* colonisation of conventional mice is well established [13],[29], it does have limitations. Delivery of a large concentration of the colonising strain into the NT results in transient colonisation with mice clearing *S. aureus* from their NT within between 28-42 days post-colonisation [13],[29]. To supplement the limitations of this transient colonisation, data in this study was also generated in GF mice that were mono-colonised with *S. aureus* from birth. This was done to better mimic persistent long-term *S. aureus* colonisation that occurs in humans, who are often colonised from infancy, primarily due to environmental exposure [1],[45]. Findings in these mice suggest that CD4+, but not γδ+, TRM cells, can be established with *S. aureus* colonisation alone in the absence of other microbial species, and support a significant role for CD4+ T cells in providing localised protection in the NT during *K. pneumoniae* infection. With the microbiome and immune cell function being dynamically related [46], it is perhaps unsurprising that mono-colonised mice would have a more developed immune system than their GF counterparts. However, these memory CD4+ T-cells were not found systemically and were instead localised to the NT where *S. aureus* resided, agreeing with what is universally accepted about TRM cells being localised, and undergoing reactivation, at the site of activation [47],[48]. Of course there are limitations to mono-colonisation, given that in humans *S. aureus* colonisation exists against the background of a diverse microbiome. Nevertheless, this approach does allow for attribution of specific immunological responses directly to *S. aureus* exposure in the absence of confounding factors.

While data here demonstrates an expansion of CD4+ TRM cells in the *S. aureus* colonised NT, heterologous immune protection cannot be specifically attributed to this cell population alone. It is likely that multiple memory cell types are functionally influenced by colonisation with *S. aureus,* and contribute to protective immune responses, potentially working in concert. This could also include innate immune cells, given that *S. aureus* has previously been shown to induce innate immune training [49]. Delineating the contribution of innate vs adaptive immune memory to *S. aureus* induced heterologous protection requires further investigation.

Overall, this study establishes that *S. aureus* colonisation expands TRM cells within the NT which may have wide-reaching benefits for the host, including enhanced protection during heterologous infection.

## Materials & methods

### Ethics statement

Animal experiments were conducted in accordance with the recommendations and guidelines of the Health Products Regulatory Authority (HPRA), the competent authority in Ireland, and in accordance with protocols approved by Trinity College Dublin (TCD) Animal Research Ethics Committee.

### Bacterial strains and growth conditions

Stocks of bacterial strains were prepared in 70% glycerol and stored at −80°C. *S. aureus* strains were grown from frozen stocks on TSA plates. SmR *S. aureus* strains were grown from frozen stocks on TSA plates supplemented with 0.001% streptomycin. Overnight stock plates were then used to prepare bacterial inoculum. *K. pneumoniae* strains were grown from frozen stocks on Luria broth (LB) agar plates. Agar plates were incubated at 37°C overnight. One *K. pneumoniae* colony from the overnight culture was cultured in LB broth in a shaking incubator overnight at 37°C. A 1:10 dilution of the overnight culture was refreshed in LB broth for a further two and a half hours prior to preparation of the bacterial inoculum.

### Preparation of bacterial inoculum

*S. aureus* bacterial colonies were collected from stock plates using an inoculating loop and resuspended in 1ml of sterile PBS to create a homogenous solution. All bacterial suspensions were measured at an optical density (OD) of 600nm. The OD_600_ was adjusted to the desired concentration, where an OD_600_ of 1 corresponds to 1×10^9^ *S. aureus* CFU/ml or 5×10^8^ *K. pneumoniae* CFU/ml. Serial dilutions of the working inoculum were plated on TSA plates or LB plates respectively for overnight growth at 37°C. CFUs were counted the next day to confirm inoculum concentrations.

### Preparation of heat-killed bacteria stocks

A single bacterial colony was transferred from an overnight stock plate to 15ml of broth, which was placed in a 37°C cell shaker overnight. The next day the broth was centrifuged at 3,000 rotations per minute (rpm) for 5 minutes, and bacteria was resuspended in PBS to form a homogenous solution. The OD_600_ was adjusted to an OD of 1 and the stock was heated at 90°C for 45 minutes. 100µl of stock was plated on TSA plates to confirm heat-killing.

### Mice

Conventional wild-type C57BL/6JCrl mice were obtained from Charles River Laboratory or bred in-house under specific pathogen free (SPF) conditions in the TCD Comparative Medicine Unit (CMU). Gnotobiotic mice (C57/BlkNTac background) were bred in-house under GF conditions in aseptic rigid walled containment isolators in the CMU. Inclusion criteria for experiments were healthy mice aged 6-12 weeks old at the initiation of experiments. Both male and female mice were used, and mice were matched for age and sex in all experiments.

### In vivo administration of Streptomycin

A 500mg/ml solution of the antibiotic streptomycin was prepared by diluting 5g of streptomycin powder in 10ml of sterile water. This stock solution was divided into 1ml aliquots which were stored at −20°C and defrosted as needed. 300µl of 500mg/ml streptomycin was administered in 300ml of drinking water to a final concentration of 0.5mg/ml for the duration of i.n. inoculation experiments. Water was refreshed every 5-7 days. Streptomycin drinking water was administered for the duration of experiments where conventional mice were colonised with *S. aureus but* was replaced with non-antibiotic treated water 2 days prior to infection in *K. pneumoniae* challenge experiments.

### In vivo administration of FTY720

A 25mg/ml solution of FTY720 was prepared by reconstituting one 25mg vial in 1ml of de-ionised water. This solution was then aliquoted, stored at −20°C and defrosted as needed. The average weight of the mice in a cage was calculated and FTY720 was administered in drinking water at a concentration of 0.3mg/kg weight/day, where 7ml was taken as the average daily water intake [18]. This water was refreshed every 5 days.

### In vivo model of nasal colonisation

SPF mice were sedated using isoflurane prior to i.n. inoculation. GF mice were fully conscious for i.n. inoculation. Unpublished data from the lab has demonstrated that the use of isoflurane has no effect on colonisation levels by *S. aureus*. *S. aureus* strain Newman SmR was administered at a concentration of 2×10^8^ CFU/per nostril in 10μl. Mice exhibit no clinical symptoms of infection, and previous studies have shown limited detection of *S. aureus* in the lung using this colonisation method [42]. *K. pneumoniae* strain ATCC Kp43816 was administered at a concentration of 2.5×10^3^ CFU/ per nostril in 25μl, which was sufficient to drive non-lethal infection. To establish long-term colonisation in germ free mice, *S. aureus* strain Newman was introduced into the bedding of mice in a sterile isolator under negative pressure. Colonisation was monitored through the collection of faecal samples at regular intervals for CFU enumeration. Colonised mice could be housed for up to 6 months and were used for breeding to generate offspring that were mono-colonised with *S. aureus* from birth. In all experiments GF mice, or offspring of *S. aureus* mono-colonised mice that were colonised from birth, were used from ages 6-12 weeks old.

### Intravenous injection with CD45

Mice were warmed to 37°C in a heating box for 10 minutes prior to injection in order to induce vasodilation of the tail vein. Mice were restrained in a restraining tube which allowed access to the tail, and administered 10µl of CD45 fluorochrome-labelled antibody, which was diluted to a final volume of 200µl in PBS. Following injection, pressure was placed on the injection site to prevent bleeding, and the mouse was returned to their cage for 10 minutes, after which they were sacrificed.

### In vivo PAMP or antibody treatment

Mice were sedated using isoflurane prior to i.n. administration of immunomodulatory molecules: *E. coli* derived LPS was i.n. administered (10μg/nostril in 10μl). Anti-IL-17 mAbs or relevant isotype control Ab was i.n. administered (50µg/nostril in 10μl) at the same time as bacterial colonisation, as well as a second i.n. administration (50µg/nostril in 10μl) 24 hours later.

### Isolation of Tissues

The area surrounding the nose was cleaned using sterile ethanol wipes and the nose was excised (approximately 1cm deep). To extract the NT, the lower jaw was first removed to access the Nasal Associated Lymphoid Tissue (NALT), which is the ridged palate at the roof of the mouth. This tissue was extracted using a scalpel and peeled away using forceps. Small tweezers were then used to scrape out and remove the tissue beneath this area. To remove the lung, liver, kidney or spleen, 70% ethanol was used to clean the mouse’s abdomen, the skin was incised, and tissue was removed.

### Isolation of cells

NT, lungs, spleens and LNs were transferred to sterile RPMI media. Lungs were transferred from their collection tube to the lid of a sterile petri dish, where they were cut into small pieces using scissors. Both chopped lung tissue samples and NT samples were digested in 1ml of digestion mix DNase (1mg/ml; Applichem Lifescience) and Collagenase D (1mg/ml; Sigma Aldrich) and placed in a 37°C cell shaker for 50 and 40 minutes respectively. Following digestion, tubes were removed from the cell shaker and diluted with RPMI media to stop enzyme activity. Spleens and LNs were mechanically digested. Tissue was then passed through either a 40µm (NT) or 70µm (Lung, spleen, LNs) cell strainers and centrifuged. Red blood cell lysis was performed using ACK lysis buffer and cells were centrifuged and resuspended in RPMI media for Trypan Blue cell counting.

### Assessment of tissue bacterial burden

Once harvested, the nose, lungs, kidney, and liver were each placed in 1ml of sterile PBS. The tissue was homogenised using either a blade homogenizer, (Kinematica Polytron PT 2500E), cleaning the blade between samples using 1M NaOH, sterile H_2_O, 70% (v/v) EtOH and sterile PBS, or a bead homogeniser (Benchmark BeadBug 6 Six Position Homogenizer 230V), where tissue samples and PBS are placed in pre-filled tubes with sterilised zirconium beads. Assessment of bacterial burden in the NT was performed on supernatant from enzymatically digested NT, as described previously [50] after initial centrifuge step and before red blood cell lysis. For enumeration of *S. aureus* bacterial burden, 20µl of homogenate was dropped in duplicate onto streptomycin-supplemented TSA plates at a range of dilutions. For enumeration of *K. pneumoniae* bacterial burden, 20µl of homogenate was dropped in duplicate onto LB plates at a range of dilutions. Plates were incubated overnight at 37°C. The number of colonies visible on the plates were counted, and CFU/ml was determined.

### Antigen recall assay

Whole isolated NT cells were cultured in 96-well plates at a concentration of 2.5×10^6^cells/ml, in a final volume of 200µl/well. Cells were stimulated with heat killed (HK)-SA (1μg/ml) or HK-KP (1μg/ml) in combination with αCD28 and αCD49d. Control wells were treated with media alone (negative control), or plate bound αCD3 and soluble αCD28 (positive control). CD4+ NT cells were isolated using Miltenyi Biotec CD4 (L3T4) positive selection microbeads as per manufacturer’s instructions and plated at 2.5×10^5^cells/ml, at a final volume of 200µl/well. CD4+ NT cells were stimulated with HK bacteria, predetermined combinations of recombinant murine cytokines IL-1β (MSC), IL-23 (MSC), IL-18 (MSC), IL-7 (Bio-Techne) and IL-12 (Bio-Techne) all at 10ng/ml, or supernatants from bacterial infected BMDCs (1:1 dilution in cell media). Control wells were treated with media alone, or plate bound αCD3 and soluble αCD28. In some experimental setups, NT CD4+ T-cells were cultured in the presence of APCs comprising total irradiated splenocytes. Spleens were isolated from naive mice and filtered through 70µm cell strainers. Cells were centrifuged and red-blood cell lysis was performed. Cells were then irradiated using a GammaCell 3000 137 Caesium Irradiator (40Gy for 7 minutes). Irradiated splenocytes were cultured with CD4+ T-cells at a ratio of 1:1. Cells were cultured at 37°C and supernatants were harvested 72 hours later for quantification of cytokine concentrations by Enzyme-Linked Immunosorbent Assay (ELISA) as per R&D kit specifications.

### BMDC cell culture

To isolate bone marrow cells, bones were flushed with cRPMI media. Cells were strained through a 70μm filter before being centrifuged at 300g for 5 minutes. Red blood cell lysis was performed using ACK lysis buffer and cells are centrifuged and resuspended in cRPMI media for Trypan Blue cell counting. Cells were plated at a concentration of 6.25×10^5^cells/ml in petri dishes in 10ml cRPMI containing 20ng/ml GM-CSF and cultured at 37°C. On day 3, cells were supplemented with 10ml of cRPMI containing 20ng/ml GM-CSF. On day 6 media was removed from cells to remove any non-adherent cells and 10ml of fresh cRPMI containing 20ng/ml GM-CSF was added to each plate. On day 8 cells were again supplemented with 10ml of cRPMI containing 20ng/ml GM-CSF. On day 10 semi-adherent cells were removed by gently pipetting the media in the dishes, centrifuged and resuspended in cRPMI media for Trypan Blue cell counting. Cells were plated at 1×10^6^ cells/ml in 96-well round-bottom plates and rested for 3 hours before stimulation. BMDCs were stimulated with live *K. pneumoniae* (MOI 50) or medium alone for one hour, after which infected media was substituted for media supplemented with gentamicin (100μg/ml, Sigma Aldrich) for 23 hours. Supernatants were used to stimulate NT CD4+ T-cells in vitro.

### Flow cytometry

Isolated NT cells were stimulated with PMA (10ng/ml; Sigma Aldrich), ionomycin (500ng/ml; Sigma Aldrich) and BFA (5µg/ml; Sigma Aldrich), for 4 hours at 37°C prior to intracellular cytokine staining for flow cytometric analysis. Cells were then washed in PBS, stained with Fixable Viability Dye-eFluor506 (diluted 1:1000 in PBS) and incubated at 4°C for 15 minutes. Cells were then washed in MACS buffer (0.5% (w/v) BSA 2mM EDTA (Sigma Aldrich) 1X PBS) followed by incubation in Fc block (αCD16/CD32, ThermoFisher) (diluted 1:100 in MACS buffer). A master mix was prepared of fluorochrome-conjugated mAbs specific for T-cell surface markers: CD45-BV570 (30-F11; Biolegend), CD3-Spark Blue 55 (M5/114.15.2; Biolegend), CD4-BB700 (RM4-5; BD Horizon), CD44-BV711 (IM7; BD Horizon), CD44-PEeFlour 610 (IM7; Invitrogen), CD69-APC-Cy7 (H1.2F3; Biolegend), CD103-BV711 (M290; MSC), CD8a-Pacific Blue (5H10; Invitrogen), or neutrophil cell surface markers; F480-ef450 (BM8; Invitrogen), Ly6G-BV650 (1A8-Biolegend), CD11c-APC-ef780 (N418; Invitrogen), CD11b-APC (C67F154-Invitrogen) and MHCII-PE-cy7 (M5/114.15.2; Invitrogen) which were diluted between 1:50 and 1:200 in MACS buffer as previously determined by titration assays. Cell samples were suspended in 50µl mAb master mix and incubated for 15 minutes at 4°C. 50µl of Fix and Perm Solution A was added to each sample to fix cells and incubated at room temperature for 15 minutes. Cells were washed twice in MACS buffer. Samples were incubated with Forkhead box protein 3 (Foxp3) Fixation/Permeabilization working solution (Fixation/Permeabilization Concentrate diluted 1:4 with Fixation/Permeabilization Diluent) at 4°C overnight. Cells were then washed twice with 1X Permeabilization buffer (10X Permeabilization buffer diluted 1:10 in dH2O). A master mix was prepared of fluorochrome-conjugated mAbs specific for intracellular and intra-nuclear proteins; (Ki67-AF700 (SolA15; Invitrogen), IL-17-BV785 (TC11-18H10.1; Biolegend) which were diluted either 1:100 or 1:50 in 1X Permeabilization buffer, as previously determined by titration assays, and incubated in the dark at 4°C for 1 hour. Cells were then washed once with 1X Permeabilization Buffer, followed by one wash with MACS buffer. Samples were re-suspended in 200µl of MACS buffer and stored in the dark at 4°C until analysis. Flow cytometry data was acquired using LSRFortessa (BD) with FACS DIVA software or Aurora with Cytek Biosciences software. FlowJo software (Treestar, Inc.) was used for post-acquisition analysis. Gating was set on respective Fluorescence Minus One (FMO) samples.

### Statistics and Reproducibility

Statistical analyses were performed using GraphPad Prism 10 software (GraphPad Software, La Jolla, CA). Differences between two groups were analysed using an unpaired Student t test. Comparisons between 3 or more groups were performed by one-way Analysis of variance (ANOVA) with a Tukey comparison post-test to compare all means, or Dunnett’s post-test if means were compared to a predetermined control group. Analysis when 2 or more variables exist were performed by two-way ANOVA with a Tukey comparison post-test to compare all means, or Sidak’s post-test to compare a subset of means. P values less than 0.05 were considered significant.

## CRediT authorship contribution statement

**Clíodhna Daly**: Writing – review & editing, Writing – original draft, Methodology, Investigation, Formal analysis, Data curation, Conceptualisation. **Seán Cahill**: Writing – review, Investigation, Formal analysis, Data curation. **Charlotte Leane:** Writing – review, Investigation, Data curation. **Simon Carlile**: Writing - review, Investigation. **Alanna Kelly**: Writing – review, Investigation. **Jenny Mannion:** Writing – review, Investigation. **Rachel McLoughlin**: Writing – review & editing, Validation, Supervision, Project administration, Methodology, Investigation, Funding acquisition, Conceptualisation.

## Declaration of competing interest

The authors have no competing interests to declare.

## Data availability

Source Data are provided with this paper and all data available on request to the corresponding author.

## Funding

This work was supported by a Wellcome Investigator Award (202846/Z/16/Z) and a Research Ireland Frontiers for the Future Award (23/-FFPA/12244) to R.M.M.

## Acknowledgements

Thank you to Dr Barry Moran and the TCD flow core, as well as Dr Stephen Lalor for advice and guidance with germ-free experiments

---

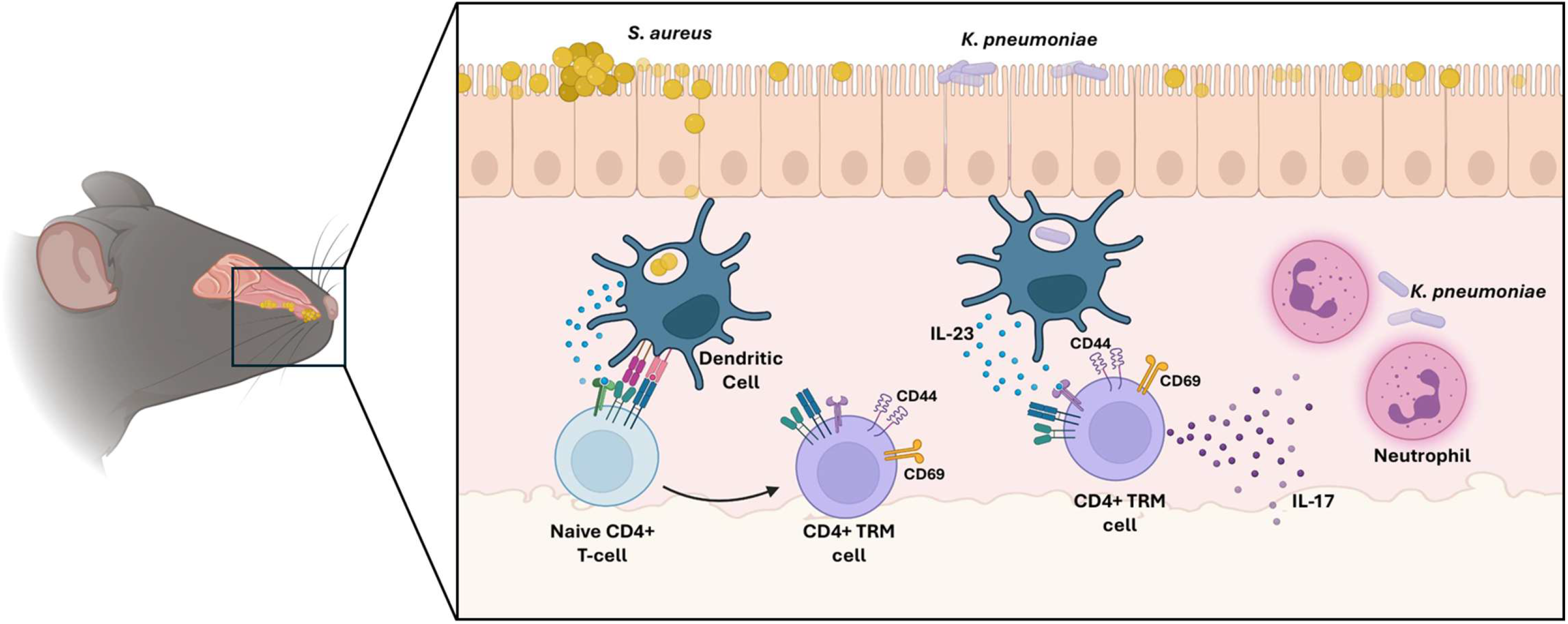

**CD4+ TRM cells are primed by *S. aureus* exposure for enhanced heterologous immune protection**

Intranasal inoculation of of wild-type C57BL6/JCrl mice with *S. aureus* strain Newman results in expansion of CD4+ TRM cells in the colonised nasal tissue. CD4+ TRM cells exhibit enhanced re-activation capacity in response to subsequent *S. aureus* exposure or Th17 polarising cytokines. IL-17 production by CD4+ TRM cells enhances protection against invading pathogens through neutrophil recruitment.

## Notes

### Competing Interest Statement

The authors have declared no competing interest.

## References

1. Sakr A, Brégeon F, Mège JL, Rolain JM, Blin O. Staphylococcus aureus Nasal Colonization: An Update on Mechanisms, Epidemiology, Risk Factors, and Subsequent Infections. Front. Microbiol. 2018; 9. Available at: /pmc/articles/PMC6186810/ [Accessed May 12, 2022].DOI: 10.3389/FMICB.2018.02419.

2. Paul O Verhoeven, Julie Gagnaire, Elisabeth Botelho-Nevers, Florence Grattard, Anne Carricajo, Frédéric Lucht, Bruno Pozzetto, et al. Detection and clinical relevance ofStaphylococcus aureusnasal carriage: an update. Expert Review of Anti-Infective Therapy, 12(1), 75–89 | 10.1586/14787210.2014.859985. 2013. Available at: https://sci-hub.se/10.1586/14787210.2014.859985 [Accessed October 9, 2024].

3. Ikuta KS, Swetschinski LR, Aguilar GR, Sharara F, Mestrovic T, Gray AP, Weaver ND, et al. Global mortality associated with 33 bacterial pathogens in 2019: a systematic analysis for the Global Burden of Disease Study 2019. Lancet. 2022; 400:2221. Available at: https://pmc.ncbi.nlm.nih.gov/articles/PMC9763654/ [Accessed December 18, 2024].DOI: 10.1016/S0140-6736(22)02185-7.

4. Pivard M, Moreau K, Vandenesch F. Staphylococcus aureus Arsenal To Conquer the Lower Respiratory Tract. mSphere. 2021; 6. Available at: https://pubmed-ncbi-nlm-nih-gov.elib.tcd.ie/34011681/ [Accessed February 22, 2022].DOI: 10.1128/MSPHERE.00059-21.

5. Guo Y, Song G, Sun M, Wang J, Wang Y. Prevalence and Therapies of Antibiotic-Resistance in Staphylococcus aureus. Front. Cell. Infect. Microbiol. 2020; 10. Available at: /pmc/articles/PMC7089872/ [Accessed May 14, 2022].DOI: 10.3389/FCIMB.2020.00107.

6. Cahill SC, Soldaini E, McLoughlin RM. Pathobiont immune tuning: the untold consequences of Staphylococcus aureus exposure on host immunity. Trends Microbiol. 2025. Available at: https://www.sciencedirect.com/science/article/pii/S0966842X25002197?via%3Dihub [Accessed September 8, 2025].DOI: 10.1016/J.TIM.2025.07.008.

7. Montgomery CP, Daniels M, Zhao F, Alegre ML, Chong AS, Daum RS. Protective Immunity against Recurrent Staphylococcus aureus Skin Infection Requires Antibody and Interleukin-17A. Infect. Immun. 2014; 82:2125. Available at: https://pmc.ncbi.nlm.nih.gov/articles/PMC3993461/ [Accessed December 9, 2024].DOI: 10.1128/IAI.01491-14.

8. Kelly AM, McCarthy KN, Claxton TJ, Carlile SR, Brien ECO, Vozza EG, Mills KHG, et al. IL-10 inhibition during immunization improves vaccine-induced protection against Staphylococcus aureus infection. JCI Insight. 2024; 9. Available at: /pmc/articles/PMC11383370/ [Accessed October 10, 2024].DOI: 10.1172/JCI.INSIGHT.178216.

9. Tsai CM, Caldera JR, Hajam IA, Chiang AWT, Tsai CH, Li H, Díez ML, et al. Non-protective immune imprint underlies failure of Staphylococcus aureus IsdB vaccine. Cell Host Microbe. 2022; 30:1163–1172.e6.DOI: 10.1016/J.CHOM.2022.06.006.

10. L Wertheim HF, Vos MC, Ott A, van Belkum A, Voss A, J W Kluytmans JA, J van Keulen PH, et al. Risk and outcome of nosocomial Staphylococcus aureus bacteraemia in nasal carriers versus non-carriers. Lancet. 2004; 364:703–708. Available at: www.thelancet.com [Accessed May 20, 2022].DOI: 10.1016/S0140-6736(04)16897-9.

11. Archer NK, Harro JM, Shirtliff ME. Clearance of Staphylococcus aureus Nasal Carriage Is T Cell Dependent and Mediated through Interleukin-17A Expression and Neutrophil Influx. 2013.DOI: 10.1128/IAI.00084-13.

12. Archer NK, Adappa ND, Palmer JN, Cohen NA, Harro JM, Lee SK, Miller LS, et al. Interleukin-17A (IL-17A) and IL-17F Are Critical for Antimicrobial Peptide Production and Clearance of Staphylococcus aureus Nasal Colonization. 2016.DOI: 10.1128/IAI.00596-16.

13. Mulcahy ME, Leech JM, Renauld JC, Mills KHG, McLoughlin RM. Interleukin-22 regulates antimicrobial peptide expression and keratinocyte differentiation to control Staphylococcus aureus colonization of the nasal mucosa. Mucosal Immunology 2016 9:6. 2016; 9:1429–1441. Available at: https://www-nature-com.elib.tcd.ie/articles/mi201624 [Accessed April 9, 2022].DOI: 10.1038/MI.2016.24.

14. Clara Padoveze M, de Jesus Pedro R, Blum-Menezes D, José Bratfich O, Luiza Moretti M, Paulo S. Staphylococcus aureus nasal colonization in HIV outpatients: Persistent or transient?DOI: 10.1016/j.ajic.2007.05.012.

15. Masopust D, Soerens AG. Tissue-Resident T Cells and Other Resident Leukocytes. Annu. Rev. Immunol. 2019; 37:521. Available at: /pmc/articles/PMC7175802/ [Accessed March 19, 2024].DOI: 10.1146/ANNUREV-IMMUNOL-042617-053214.

16. Curham LM, Mannion JM, Daly CM, Wilk MM, Borkner L, Lalor SJ, McLoughlin RM, et al. Bystander activation of Bordetella pertussis-induced nasal tissue-resident memory CD4 T cells confers heterologous immunity to Klebsiella pneumoniae. Eur. J. Immunol. 2023; 53:2250247. Available at: https://onlinelibrary.wiley.com/doi/full/10.1002/eji.202250247 [Accessed June 8, 2023].DOI: 10.1002/EJI.202250247.

17. Bedford JG, Caminschi I, Wakim LM. Intranasal Delivery of a Chitosan-Hydrogel Vaccine Generates Nasal Tissue Resident Memory CD8 + T Cells That Are Protective against Influenza Virus Infection. Available at: www.mdpi.com/journal/vaccines [Accessed August 24, 2021].DOI: 10.3390/vaccines8040572.

18. Allen AC, Wilk MM, Misiak A, Borkner L, Murphy D, Mills KHG. Sustained protective immunity against Bordetella pertussis nasal colonization by intranasal immunization with a vaccine-adjuvant combination that induces IL-17-secreting T RM cells. Mucosal Immunol. 2018; 11:1763–1776.DOI: 10.1038/s41385-018-0080-x.

19. Wilk MM, Borkner L, Misiak A, Curham L, Allen AC, Mills KHG. Immunization with whole cell but not acellular pertussis vaccines primes CD4 T RM cells that sustain protective immunity against nasal colonization with Bordetella pertussis. 2019. Available at: 10.1080/22221751.2018.1564630 [Accessed August 24, 2021].DOI: 10.1080/22221751.2018.1564630.

20. Borkner L, Curham LM, Wilk MM, Moran B, Mills KHG. IL-17 mediates protective immunity against nasal infection with Bordetella pertussis by mobilizing neutrophils, especially Siglec-F+ neutrophils. Mucosal Immunology 2021 14:5. 2021; 14:1183–1202. Available at: https://www.nature.com/articles/s41385-021-00407-5 [Accessed August 30, 2021].DOI: 10.1038/s41385-021-00407-5.

21. O’hara JM, Redhu NS, Cheung E, Robertson NG, Patik I, Sayed S El, Thompson CM, et al. Generation of protective pneumococcal-specific nasal resident memory CD4 + T cells via parenteral immunization. 2019. Available at: 10.1038/s41385-019-0218-5 [Accessed August 27, 2021].DOI: 10.1038/s41385-019-0218-5.

22. Christo SN, Evrard M, Park SL, Gandolfo LC, Burn TN, Fonseca R, Newman DM, et al. Discrete tissue microenvironments instruct diversity in resident memory T cell function and plasticity. NATuRE IMMuNOLOGY |. 2021; 22. Available at: 10.1038/s41590-021-01004-1 [Accessed June 12, 2023].DOI: 10.1038/s41590-021-01004-1.

23. Muñoz-Ruiz M, Llorian M, D’Antuono R, Pavlova A, Mavrigiannaki AM, McKenzie D, García-Cassani B, et al. IFN-γ–dependent interactions between tissue-intrinsic γδ T cells and tissue-infiltrating CD8 T cells limit allergic contact dermatitis. Journal of Allergy and Clinical Immunology. 2023; 152:1520–1540. Available at: https://www.sciencedirect.com/science/article/pii/S0091674923009818?via%3Dihub [Accessed April 29, 2025].DOI: 10.1016/J.JACI.2023.07.015.

24. Naik S, Bouladoux N, Linehan JL, Han SJ, Harrison OJ, Wilhelm C, Conlan S, et al. Commensal–dendritic-cell interaction specifies a unique protective skin immune signature. Nature 2015 520:7545. 2015; 520:104–108. Available at: https://www-nature-com.elib.tcd.ie/articles/nature14052 [Accessed June 2, 2022].DOI: 10.1038/NATURE14052.

25. Nan K, Zhong Z, Yue Y, Shen Y, Zhang H, Wang Z, Zhuma K, et al. Fasting-mimicking diet-enriched Bifidobacterium pseudolongum suppresses colorectal cancer by inducing memory CD8+ T cells. Gut. 2025; 74:775–786. Available at: https://gut.bmj.com/content/74/5/775 [Accessed July 29, 2025].DOI: 10.1136/GUTJNL-2024-333020.

26. Anderson KG, Mayer-Barber K, Sung H, Beura L, James BR, Taylor JJ, Qunaj L, et al. Intravascular staining for discrimination of vascular and tissue leukocytes. Nat. Protoc. 2014; 9:209. Available at: /pmc/articles/PMC4428344/ [Accessed April 6, 2022].DOI: 10.1038/NPROT.2014.005.

27. Brinkmann V, Cyster JG, Hla T. FTY720: Sphingosine 1-Phosphate Receptor-1 in the Control of Lymphocyte Egress and Endothelial Barrier Function. American Journal of Transplantation. 2004; 4:1019–1025. Available at: https://onlinelibrary.wiley.com/doi/full/10.1111/j.1600-6143.2004.00476.x [Accessed April 21, 2022].DOI: 10.1111/J.1600-6143.2004.00476.X.

28. Hofmann M, Brinkmann V, Zerwes HG. FTY720 preferentially depletes naive T cells from peripheral and lymphoid organs. Int. Immunopharmacol. 2006; 6:1902–1910. Available at: https://pubmed.ncbi.nlm.nih.gov/17161343/ [Accessed October 14, 2024].DOI: 10.1016/J.INTIMP.2006.07.030.

29. Kiser KB, Cantey-Kiser JM, Lee JC. Development and Characterization of a Staphylococcus aureus Nasal Colonization Model in Mice. Infect. Immun. 1999; 67:5001–5006.

30. Maschirow L, Suttorp N, Opitz B. Microbiota-dependent regulation of antimicrobial immunity in the lung. Am. J. Respir. Cell Mol. Biol. 2019; 61:284–289. Available at: www.atsjournals.org [Accessed June 9, 2022].DOI: 10.1165/RCMB.2019-0101TR/SUPPL_FILE/DISCLOSURES.PDF.

31. Yu S, Wang K, Cao C, Zhang B, Chen Y, Wu C, Li C, et al. Tissue-resident memory T cells exhibit phenotypically and functionally heterogeneous in human physiological and pathological nasal mucosa. Clinical Immunology. 2024; 258:109860.DOI: 10.1016/J.CLIM.2023.109860.

32. Mckinstry KK, Mills KHG, Wilk MM. CD4 T RM Cells Following infection and immunization: implications for More effective vaccine Design. 2018; 9:1860. Available at: www.frontiersin.org. DOI: 10.3389/fimmu.2018.01860.

33. Carolina M, Vesely A, Pallis P, Bielecki P, Low JS, Zhao J, Harman CCD, et al. Effector T H 17 cells give rise to long-lived T RM cells that are essential for an immediate response against bacterial infection.DOI: 10.1016/j.cell.2019.07.032.

34. Schmitt P, Borkner L, Jazayeri SD, McCarthy KN, Mills KH. Nasal vaccines for pertussis. Curr. Opin. Immunol. 2023; 84:102355.DOI: 10.1016/J.COI.2023.102355.

35. Jin C, Lagoudas GK, Zhao C, Bullman S, Bhutkar A, Hu B, Ameh S, et al. Commensal Microbiota Promote Lung Cancer Development via γδ T Cells. Cell. 2019; 176:998–1013.e16. Available at: https://pubmed.ncbi.nlm.nih.gov/30712876/ [Accessed October 29, 2024].DOI: 10.1016/J.CELL.2018.12.040.

36. Yang Q, Li C, Wang W, Zheng R, Huang X, Deng H, Jin P, et al. Infiltration pattern of gammadelta T cells and its association with local inflammatory response in the nasal mucosa of patients with allergic rhinitis. Int. Forum Allergy Rhinol. 2019; 9:1318–1326. Available at: /doi/pdf/10.1002/alr.22421 [Accessed April 28, 2026].DOI: 10.1002/ALR.22421;WEBSITE:WEBSITE:PERICLES;JOURNAL:JOURNAL:20426984;ISSUE:ISSUE:DOI.

37. Xiong H, Carter RA, Leiner IM, Tang Y-W, Chen L, Kreiswirth BN, Pamer EG. Distinct Contributions of Neutrophils and CCR2 Monocytes to Pulmonary Clearance of Different Klebsiella pneumoniae Strains. 2015. Available at: https://journals.asm.org/journal/iai [Accessed October 15, 2024].DOI: 10.1128/IAI.00678-15.

38. Mannion JM, Segal BM, McLoughlin RM, Lalor SJ. Respiratory tract Moraxella catarrhalis and Klebsiella pneumoniae can promote pathogenicity of myelin-reactive Th17 cells. Mucosal Immunol. 2023; 16:399–407. Available at: http://www.mucosalimmunology.org/article/S1933021923000296/fulltext [Accessed December 6, 2024].DOI: 10.1016/j.mucimm.2023.04.003.

39. Lee HG, Cho MZ, Choi JM. Bystander CD4+ T cells: crossroads between innate and adaptive immunity. Exp. Mol. Med. 2020; 52:1255. Available at: /pmc/articles/PMC8080565/ [Accessed October 12, 2022].DOI: 10.1038/S12276-020-00486-7.

40. Harrison E, Aggarwal D, Bellis K, Blane B, de Goffau M, Wagner J, Ng D, et al. The nasal microbiome redefines Staphylococcus aureus colonisation. 2025. Available at: https://www.researchsquare.com [Accessed September 15, 2025].DOI: 10.21203/RS.3.RS-6079410/V1.

41. Aggarwal D, Bellis KL, Blane B, de Goffau MC, Wagner J, Ng DYK, Raven KE, et al. Large-scale characterisation of the nasal microbiome redefines Staphylococcus aureus colonisation status. Nature Communications 2025 16:1. 2025; 16:10415-. Available at: https://www.nature.com/articles/s41467-025-66564-4 [Accessed June 3, 2026].DOI: 10.1038/s41467-025-66564-4.

42. Kelly AM, Leech JM, Doyle SL, McLoughlin RM. Staphylococcus aureus-induced immunosuppression mediated by IL-10 and IL-27 facilitates nasal colonisation. PLoS Pathog. 2022; 18. Available at: /pmc/articles/PMC9282462/ [Accessed October 9, 2024].DOI: 10.1371/JOURNAL.PPAT.1010647.

43. Mcloughlin RM, Dempsey DJ, Tracey ·, Claxton J, Browne E, Connolly R, Ng J, et al. Circulating T-cell phenotypes are influenced by prior Staphylococcus aureus exposure in hemodialysis patients. Available at: 10.1007/s10096-026-05581-7 [Accessed August 5, 2026].DOI: 10.1007/s10096-026-05581-7.

44. Shin H, Iwasaki A. A vaccine strategy that protects against genital herpes by establishing local memory T cells. Nature 2012 491:7424. 2012; 491:463–467. Available at: https://www.nature.com/articles/nature11522 [Accessed January 3, 2025].DOI: 10.1038/nature11522.

45. Peacock SJ, Justice A, Griffiths D, De Silva GDI, Kantzanou MN, Crook D, Sleeman K, et al. Determinants of Acquisition and Carriage of Staphylococcus aureus in Infancy. J. Clin. Microbiol. 2003; 41:5718–5725. Available at: http://mlst.net/ [Accessed May 23, 2022].DOI: 10.1128/JCM.41.12.5718-5725.2003.

46. Yuruker O, Yılmaz İ, Güvenir M. The Symbiotic Defence: Lung Microbiota and The Local Immune System.

47. Iijima N. The emerging role of effector functions exerted by tissue-resident memory T cells. 2024. Available at: 10.1093/oxfimm/iqae006 [Accessed October 17, 2024].DOI: 10.1093/oxfimm/iqae006.

48. Rotrosen E, Kupper TS. Assessing the generation of tissue resident memory T cells by vaccines. Nat. Rev. Immunol. 2023; 23:1. Available at: https://pmc.ncbi.nlm.nih.gov/articles/PMC10064963/ [Accessed November 4, 2024].DOI: 10.1038/S41577-023-00853-1.

49. Carlile SR, Cahill SC, O’Brien EC, Neto NGB, Monaghan MG, McLoughlin RM. Staphylococcus aureus induced trained immunity in macrophages confers heterologous protection against gram-negative bacterial infection. iScience. 2024; 27:111284. Available at: https://www.cell.com/action/showFullText?pii=S2589004224025094 [Accessed August 13, 2026].DOI: 10.1016/j.isci.2024.111284.

50. Jazayeri SD, Borkner L, Sutton CE, Mills KHG. Respiratory immunization using antibiotic-inactivated Bordetella pertussis confers T cell-mediated protection against nasal infection in mice. Nat. Microbiol. 2025; 10:3094. Available at: https://pmc.ncbi.nlm.nih.gov/articles/PMC12669047/ [Accessed July 3, 2026].DOI: 10.1038/S41564-025-02166-6.

