## Supplemental for "IL-17 producing tissue-resident memory T-cells expanded during *Staphylococcus aureus* nasal colonisation provide heterologous immune protection"

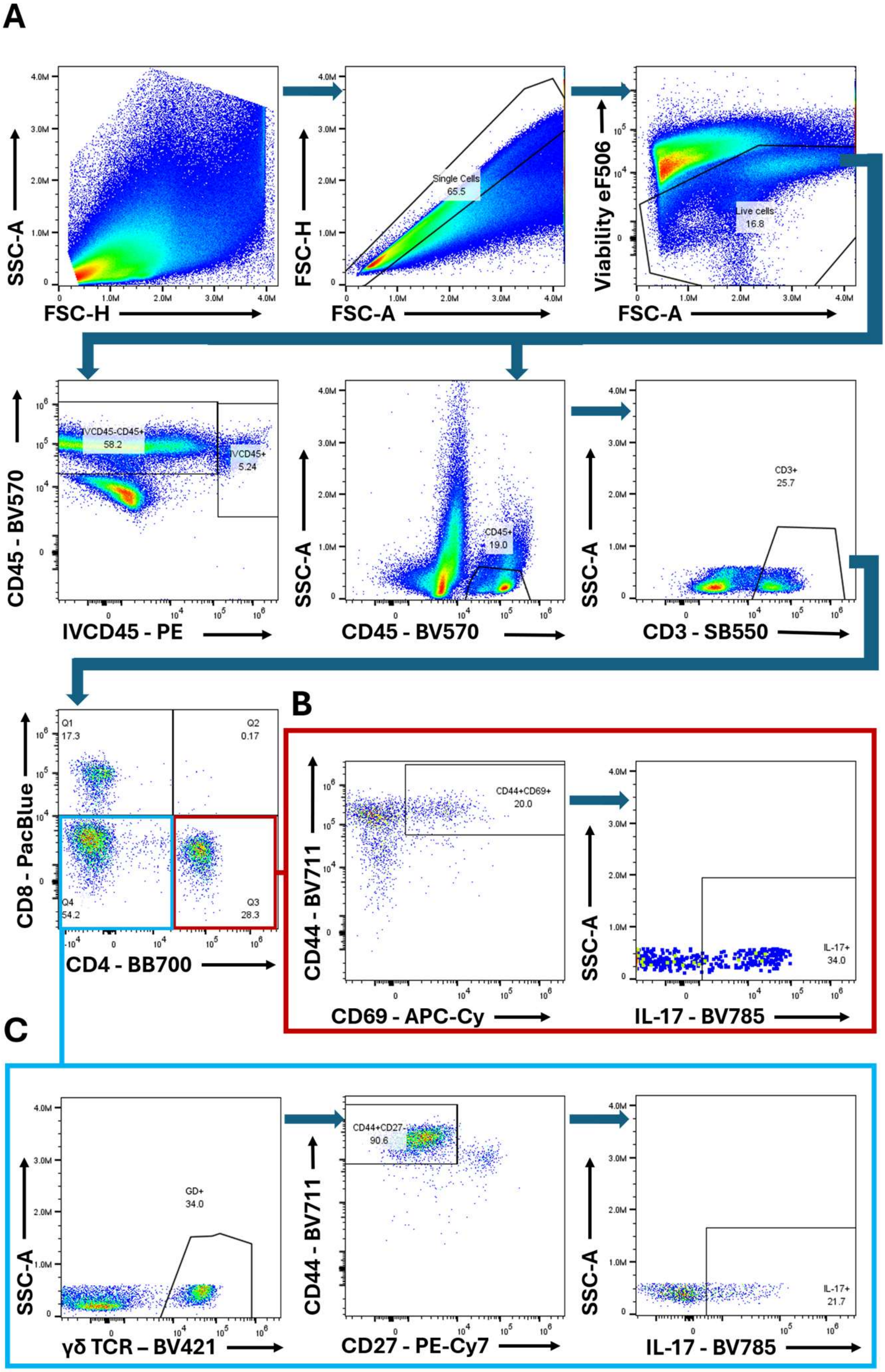


**Supplemental figure 1**

**Nasal tissue, spleen and lymph node T-cell flow cytometry gating strategy**

Plots feature cells from the NT of a *S. aureus* previously exposed mice. Cell gates were set using fluorescence minus one (FMO) samples on lymphocytes, single cells, Live cells, IVCD45-CD45+ or CD45+ cells that were CD3+ (A). Analysis of CD4+ TRM cells required cell gating on CD4+, CD44High, CD69+, CD62L- cells +/- IL-17 (B). Analysis of Resident γδ+ T cells required cell gating on CD4-, γδ T cell receptor+, CD44+CD27- cells +/- IL-17 (C).

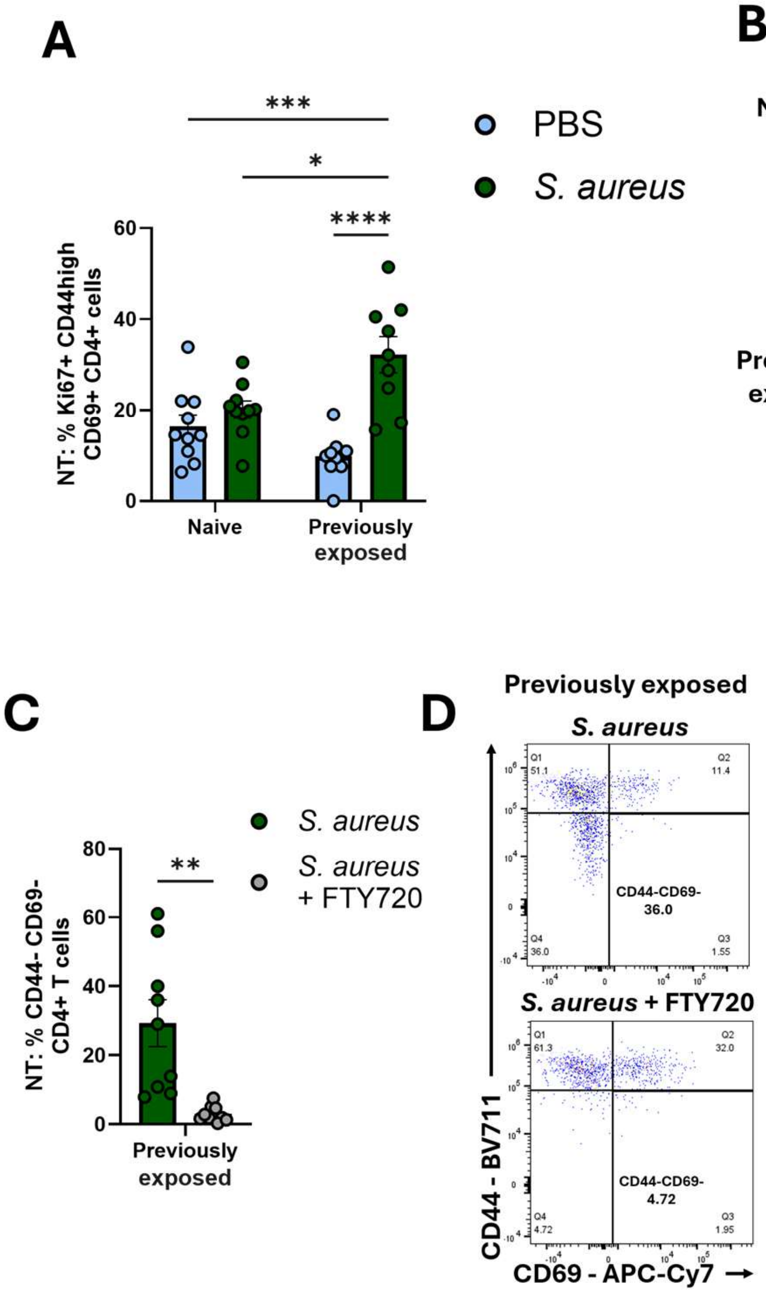


**Supplemental figure 2**

**TRM cell re-activation, proliferation and bacterial clearance in the *S. aurues* colonised NT.**

Wild-type C57BL6/JCrl mice were i.n. inoculated with *S. aureus* (Newman SmR 2x108 CFU/nostril in 10µl) or PBS on day 0. *S. aureus* (Newman SmR 2x108 CFU/nostril in 10µl), or PBS, was administered i.n. to previously exposed mice (28 or 56-days post-*S. aureus* exposure), or to naive control mice. One group of 56-day previously exposed mice received FTY720 in drinking water 10 days prior to secondary *S. aureus* administration and for the duration of the experiment. 3 days after secondary *S. aureus* exposure NT were removed, and tissue digested for flow cytometry analysis. The percentage (%) of CD4+ TRM cells (CD4+, CD44high, CD69+) expressing Ki67 were assessed in the NT of naive and 56-day previously exposed mice following re-exposure to *S. aureus* or PBS (A), with representative flow plots shown (B). The percentage (%) of NT CD4+ T-cells that are CD44- in 56-day convalescent mice following re-exposure to *S. aureus* +/- FTY720 administration prior to *S. aureus* re-exposure were assessed (C) with representative flow plots shown (D). Homogenized nose tissue and supernatants from digested NT were serially diluted and plated onto streptomycin-supplemented TSA plates. Plates were grown overnight and *S. aureus* CFU were enumerated on day 3 post- re-exposure. Results are expressed as Log CFU/g from the nose (E) and Log CFU/tissue from the NT (F) of naïve and 28-day previously exposed mice that were re-exposed to *S. aureus*. Results are expressed as mean ± S.E.M. (n=9-10 per group). Results are expressed as mean ± S.E.M. Statistical analysis was performed using two-way analysis of variance or student t test. *P ≤ 0.05, **p ≤ 0.01, ***p ≤ 0.001., ****p ≤ 0.0001.

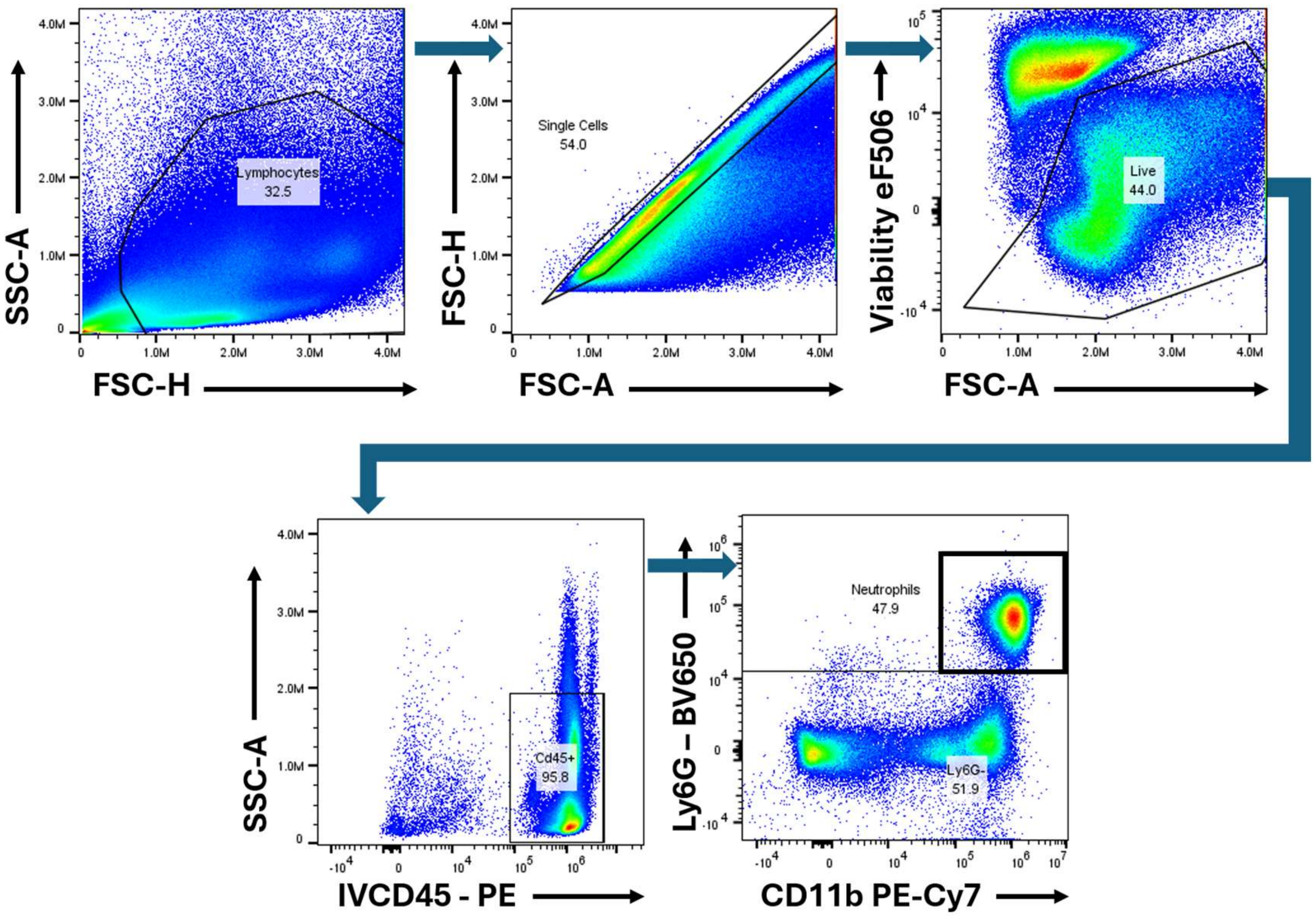


**Supplemental figure 3**

**Lung neutrophil cell flow cytometry gating strategy**

Lung cell gates were set on lymphocytes, single cells, live cells, CD45+, Ly6G+, CD11b+ cells for neutrophil analysis.

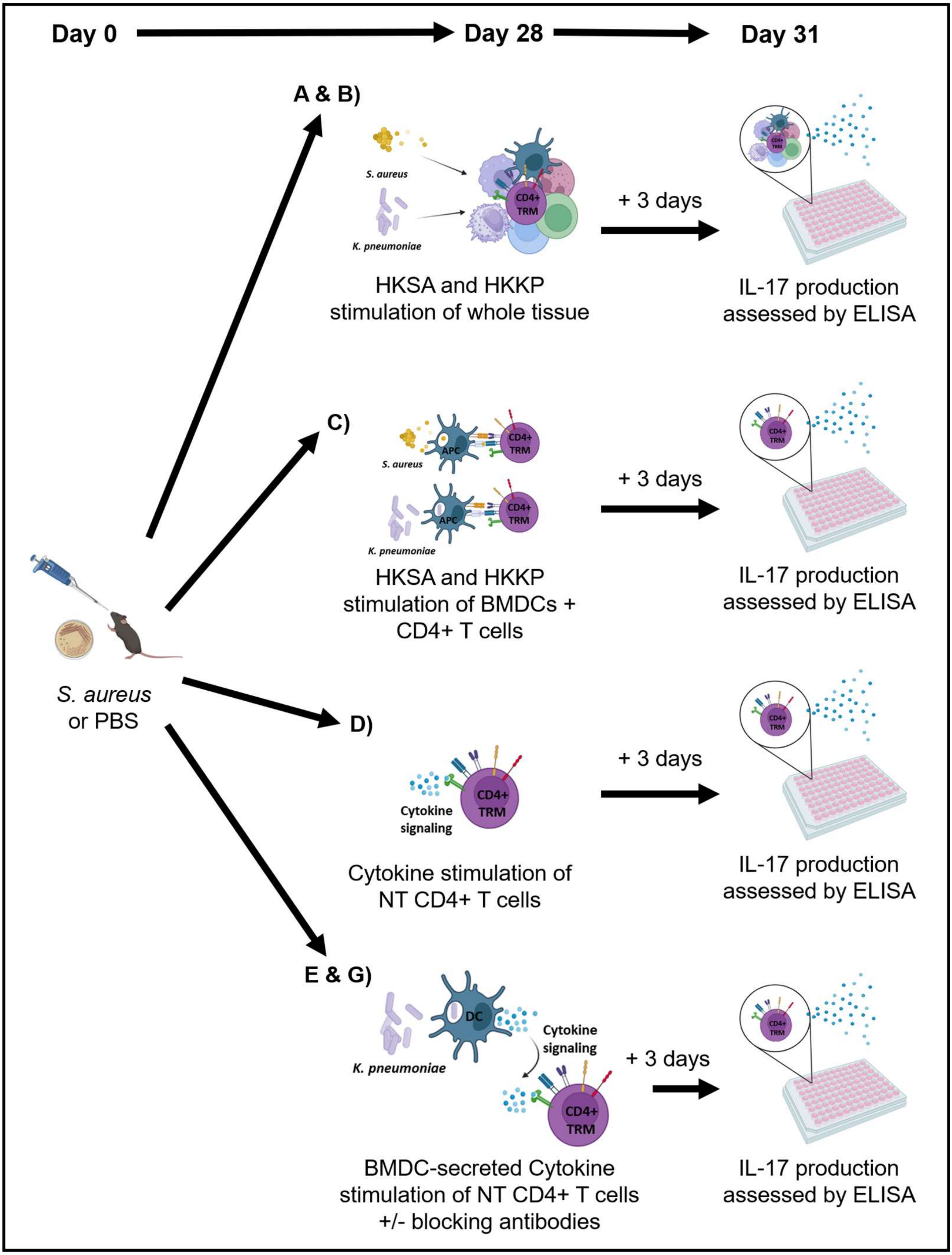


**Supplemental figure 4**

**Mechanism of action for investigating CD4+ TRM cell activation in vitro**

Wild-type C57BL6/JCrl mice were intranasally inoculated with *S. aureus* (Newman SmR $2x10^8$ CFU/nostril in 10µl) or PBS. At day 28 post-colonisation, NT was harvested and digested. Graph depicts experimental setup for data presented in Figure 5 A-G.
